# Five hundred million years of methylation: tracing the mutational origins of vertebrate genome composition

**DOI:** 10.64898/2026.09.02.747768

**Authors:** Tania Bobbo, Wageesha Widuranga, Alessio Boattini, Pietro Liò, Cristian Taccioli

## Abstract

Methylation-associated deamination removes CpG from vertebrate genomes, but how it affects the dinucleotide profile remains unresolved. We analysed 753 vertebrate and 481 invertebrate genomes to test whether CpG loss defines a compositional axis and to identify its strongest signature. CpG depletion was the dominant axis of vertebrate dinucleotide variation. Unexpectedly, its strongest between-genome correlate was AG/CT rather than the direct mutational product TpG/CpA, which showed the expected dataset-wide mass balance but varied little among genomes. A forward-evolution model based on measured seven-nucleotide human germline substitution rates produced neither CpG depletion nor AG/CT enrichment when methylated-CpG mutability was excluded. Adding one CpG-specific mutability term, calibrated only to the mammalian CpG ratio, reproduced both features, identifying AG/CT as a second-order consequence of the context-dependent mutation network. Within genomes, CpG depletion was strongest in transposable elements and weakened with distance from them. Across vertebrates, the axis followed Amniota more closely than endothermy and was associated with an expanded GC-rich isochore compartment. A Machine Learning analysis shows that CpG depletion and AG/CT were the principal features separating vertebrates from invertebrates, in which both were markedly attenuated. Thus, a methylation-associated axis organises vertebrate dinucleotide composition, and its strongest marker is not the immediate product of CpG deamination.

## Introduction

Short-sequence counts reveal a robust regularity in genomic DNA. Within either strand of most double-stranded genomes, adenine and thymine occur at similar frequencies, as do cytosine and guanine. This pattern is known as Chargaff’s second parity rule (1–3). It is weaker or absent in genomes where the mutation rate is different between strands, such as animal mitochondrial genomes, and in many single-stranded viral genomes (4, 5). The same regularity extends to adjacent bases: a dinucleotide and its reverse complement, such as CA and TG, generally have similar frequencies (4, 6, 7), whereas a dinucleotide need not match its mirror sequence or its simple complement (8). The sixteen dinucleotides can therefore be represented by ten strand-symmetric coordinates: six reverse-complement pairs (AA/TT, AC/GT, AG/CT, CA/TG, GA/TC and CC/GG) and four self-reverse-complementary dinucleotides (AT, TA, CG and GC).

One of these coordinates is exceptional in vertebrates. CpG is the principal target of DNA methylation, and a methylated cytosine deaminates to thymine much more readily than an unmethylated cytosine. Once fixed, this mutation converts CpG to TpG on one strand and to its reverse-complement representation, CpA, on the other (9, 10). Repeated over evolutionary time, the process depletes CpG and increases TpG/CpA, producing a genome-wide mass balance (11). Vertebrates generally methylate a large fraction of the nuclear genome, including the germ line, and therefore accumulate a pervasive and heritable CpG-depletion signature. Invertebrate methylation is more heterogeneous among lineages in its genomic extent, sequence context and enzymatic machinery, and is frequently associated with transposable-element content (12–14). This contrast provides a useful control: a coordinated signature caused by pervasive CpG methylation should be strong in vertebrates and attenuated across the more heterogeneous invertebrate set.

Previous comparative studies have connected CpG methylation, dinucleotide abundance and genome architecture. Simmen identified genome-wide relationships between cytosine methylation and dinucleotide frequencies, including the expected trade-off between CpG loss and TpG/CpA gain (11). Other work associated genome size with relative CpG methylation across metazoans (15) and linked DNA methylation to transposable-element expansion and genome size (16). These observations suggest that CpG depletion is not only a local mutational scar, but may be part of a broader compositional relationship among methylation, transposable elements and genome organisation.

Here we address four linked questions. First, can vertebrate dinucleotide composition be reduced to a small number of coordinated signals after base composition is removed? Second, which dinucleotide best records differences in CpG loss among genomes, and can that relationship be generated by a measured mutation process? Third, where within genomes and across vertebrate lineages is this signal concentrated? Fourth, does the same compositional regime distinguish vertebrates from invertebrates after accounting for compositional closure and taxonomic non-independence?

The main conceptual advance is not the observation that vertebrate CpG is depleted, but the identification and mechanistic explanation of its strongest secondary signature. We combine a comparative analysis of 753 vertebrate genomes with forward evolution under human germline substitution rates resolved in their complete seven-nucleotide context (17). This design separates two questions that correlation alone cannot distinguish: which coordinates track CpG loss and which of those coordinates can arise from the context-dependent mutation network after a CpG-specific methylation term is introduced. Neighbour-dependent mutation networks can generate genome-wide compositional regularities (5, 8); here we use such a network to derive the unexpected identity of the strongest between-genome signature of CpG depletion and then map that signature across genomic compartments and vertebrate lineages. Because GC-biased gene conversion (gBGC) can raise genomic G+C by favouring the fixation of G/C alleles, we also included a simplified GC-favouring term in the forward model. This term served only as a compositional correction and was not intended as a direct test of gBGC.

## Materials and Methods

All analyses were performed in Python 3.11 using NumPy, pandas, SciPy and scikit-learn. Analysis code is available at https://github.com/tacc-code-tacclab/CGtrek.

### Genome datasets and deduplication

We analysed 753 vertebrate and 481 invertebrate nuclear reference genome assemblies retrieved from NCBI RefSeq in May 2026. One assembly was retained per species. When RefSeq contained more than one eligible assembly for a species, the most recent assembly with the lowest number of Ns or indeterminate bases was used.

### Dinucleotide counting, strand symmetry and observed-over-expected ratios

For each assembly, the four bases and sixteen dinucleotides were counted on the forward strand. Only unambiguous bases were included, and a dinucleotide was not counted if it crossed an assembly-sequence boundary or a run of ambiguous bases. Let *N* denote the number of counted bases and *S* the number of counted adjacent-base positions. Base and dinucleotide frequencies were calculated as

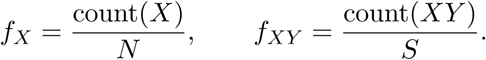

The denominator *S* was defined as the sum of the sixteen dinucleotide counts. This definition treats ambiguous runs in the same way as sequence boundaries. Using *N* minus the number of assembly sequences instead changes the denominator by up to 3.4 *×* 10*^−^*^3^ in the most fragmented assemblies. Forcing base and dinucleotide frequencies onto a common denominator multiplies all sixteen ratios within a genome by the same value, changes mean *ρ*_CG_ by less than 10*^−^*^6^ and does not alter any conclusion.

For each dinucleotide *XY*, the observed-over-expected ratio was

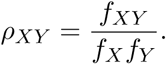

The denominator *f_X_f_Y_*is the frequency expected if the two component bases occur independently while the observed base composition of that genome is retained. Thus, *ρ* is a relative-occurrence measure rather than a raw frequency: it asks whether *XY* occurs more or less often than expected in a genome containing the same proportions of *X* and *Y* . A value of one indicates no departure from this base-composition expectation; values below and above one indicate depletion and enrichment, respectively. This normalisation allows genomes with different A, T, G and C contents to be compared without treating differences caused simply by base availability as dinucleotide-specific signals.

Intrastrand symmetry was assessed by comparing the members of each reverse-complement pair across genomes. AT, TA, CG and GC were retained individually because they are their own reverse complements. The six paired coordinates were AA/TT, AC/GT, AG/CT, CA/TG, GA/TC and CC/GG. Within each pair, the two observed frequencies and the two base-composition expectations were averaged separately, and the ratio of these averages was used. More explicitly, defining *e_XY_*= *f_X_f_Y_*and denoting the reverse complement of *XY* by rc(*XY*), each paired coordinate *d* was calculated as

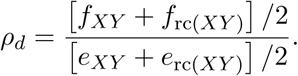

Pairing and normalisation therefore serve different purposes: pairing removes redundant strand orientation when reverse-complement parity holds, whereas *ρ* removes the contribution expected from base composition and gives the relative enrichment or depletion being compared among genomes.

The sufficiency of a strand-symmetric mutation process to generate parity was tested separately with an explicit two-strand extension of the forward model. Symmetric rates were compared with the strand-biased substitution factors measured in human mitochondrial DNA; the complete protocol and simulation outputs are provided in Supplementary Methods and Results.

### Genome-level CpG depletion and clade-block bootstrap

CpG depletion was summarised for each genome by *ρ*_CG_ and across genomes by the arithmetic mean of the genome-level ratios. We calculated confidence intervals for mean *ρ*_CG_, for its correlations with AG/CT and AC/GT, and for the TpG/CpA mass-balance statistic. These intervals used a non-parametric clade-block bootstrap. Vertebrate genomes were partitioned by NCBI taxonomic order. In each of 2000 replicates, the same number of orders as observed in the dataset was sampled with replacement, and all genomes from each selected order entered the replicate together. A repeatedly selected order contributed its complete genome set repeatedly; an unselected order contributed none. The statistic was recalculated for each replicate, and the 2.5th and 97.5th percentiles defined the 95 % confidence interval.

Taxonomic order was used as the block because it was the finest rank that provided enough units for resampling while capturing substantial between-lineage variation in *ρ*_CG_. A genome-level bootstrap would treat closely related assemblies as independent and would therefore underestimate uncertainty.

### CpG mass balance and coupling to the other coordinates

To evaluate the expected product balance of CpG deamination, we defined *u* as the CpG deficit, 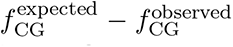, and *v* as the excess of the strand-symmetric TpG/CpA coordinate above its base-composition expectation. Because the strand-symmetric coordinate averages the two reverse-complement products, its expected excess is one half of the CpG deficit. The reported stoichiometric statistic was the ratio of the mean excess to the mean deficit, *v/u*, with theoretical expectation 0.5.

For each of the nine non-CG coordinates, we then measured how much between-genome variation was associated specifically with CpG after base composition had been included. Let *f_d_*be the observed frequency of coordinate *d* and *e_d_*its expected frequency from the component bases. We compared the nested linear models

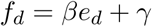

and

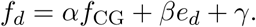

The quantity 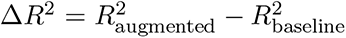 is the additional variance explained by observed CpG frequency beyond base composition. It measures association strength but not direction.

Three safeguards were applied. First, Δ*R*^2^ was recalculated under leave-one-order-out cross-validation: both models were fitted to all but one taxonomic order and evaluated on the held-out order, with every order held out once. This prevents closely related species from being divided between the training and validation sets. Second, the nested models were compared by Akaike and Bayesian information criteria, which penalise the additional parameter. Third, *p* values for the nine tested coordinates were adjusted by the Benjamini–Hochberg false-discovery-rate procedure.

We also tested two distinct sources of non-independence. Because dinucleotide frequencies form a closed composition (29), key associations were recalculated after a centred-log-ratio transformation of the complete sixteen-part dinucleotide frequency vector; paired coordinates were formed as the mean transformed value of their reverse-complement members. Because species share evolutionary history (18), associations were also estimated by generalised least squares using a three-rank taxonomic covariance: the diagonal was 1, genomes in the same order had covariance 2*/*3, genomes in the same class but different orders had covariance 1*/*3, and genomes in different classes had covariance 0. Key correlations were additionally recalculated from order and class means.

### Trek context-dependent substitution rates

The forward model used the Trek resource of human germline single-nucleotide substitution rates (17). For a central nucleotide, Trek provides its rates of substitution to each of the other three bases as a function of the complete seven-nucleotide context: the central base and the three bases on each side. The table therefore contains 4^7^ = 16,384 contexts and three possible substitutions per context, listed in alphabetical order of the destination base. Rates were inferred from substitutions accumulated in age-dated LINE-1 remnants and are expressed as relative germline substitution rates. When a seven-mer was too rare for direct estimation, Trek used the best-supported shorter context, successively falling back to a five-mer, three-mer or single base; the values analysed here already include that fallback.

The Trek core rates exclude the elevated mutability of methylated CpG, allowing that effect to be introduced as a separate calibrated term. As a quality-control check, the dominant single-base substitution in the table was the expected transition for each nucleotide: A-to-G, C-to-T, G-to-A and T-to-C.

### Forward evolution and model calibration

The principal simulation started from a random circular sequence of 2,000,000 nucleotides with A/T/G/C composition 20*/*20*/*30*/*30 %. Circularisation ensured that every position had a complete seven-nucleotide context. Each position was assigned three Trek substitution rates, and its total rate was their sum. At each step, 0.001*N* substitutions were sampled. Positions were selected in proportion to their total rates, and the destination base was selected in proportion to the three rates at that position. After applying the substitutions, rates were recalculated at each mutated position and at the three positions on either side because their seven-mer contexts had changed. Evolution continued to approximately five substitutions per site, by which point base composition was stationary, and three independent random seeds were used. Before adding any further process, the engine was required to reproduce the published Trek equilibrium A/T/G/C = 30.9*/*30.9*/*19.1*/*19.1 % (17). Additional runs from high- and low-GC starting compositions were required to converge to the same equilibrium.

CpG methylation was represented by one multiplier, *m*. The C-to-T rate was multiplied by *m* when the central cytosine was immediately followed by guanine; in the reverse-complement orientation, the G-to-A rate was multiplied by *m* when the central guanine was immediately preceded by cytosine. All other rates were unchanged. One-dimensional root finding calibrated *m* so that equilibrium *ρ*_CG_ equalled the median of the mammalian genomes. No AG/CT value was used in this calibration.

GC-biased gene conversion was approximated by a second factor, *b*, multiplying every substitution rate whose product was C or G. This factor was calibrated only so that equilibrium G+C equalled the human assembly value. The simulated and observed profiles were compared using the same reverse-complement averaging and *ρ* definitions as the genome analysis. Thus, CpG and G+C were the only fitted targets; AG/CT and the other nine coordinate values were model predictions.

### Transposable-element localisation

Genome sequences and RepeatMasker annotations were obtained from NCBI RefSeq for seventeen assemblies. Transposable elements were defined as intervals assigned to the DNA, SINE, LINE, LTR, RC or Retroposon classes. Simple repeats, low-complexity regions, satellites and RNA repeats were excluded. Overlapping and nested transposable-element intervals were merged before analysis.

For each genome, *ρ*_CG_ was calculated with region-specific base composition inside the union of transposable-element intervals, in all non-element sequence and in flanking sequence grouped by base-wise distance from the nearest element: 0–100, 100–500, 500–1000, 1000–2000, 2000–5000 and *>* 5000 bp. Non-element repeat classes were retained in the flanking sequence. As a quality-control test, genome-wide *ρ*_CG_ recalculated from each assembly was required to reproduce its value in the count table; all seventeen differed by less than 8.5*e −* 06.

Assemblies with annotated transposable-element content below 2 % were treated as incompletely annotated and excluded from pooled tests, leaving 11 genomes. Inside-element *ρ*_CG_ and the *>* 5 kb value were compared by a two-sided exact Wilcoxon signed-rank test. The association between transposable-element fraction and genome-wide *ρ*_CG_ was evaluated by partial correlation with AG/CT held constant.

### Multiple factor analysis and group-separation tests

Multiple factor analysis used two standardised variable groups: base composition (*f_A_, f_C_, f_G_, f_T_*) and the ten dinucleotide ratios. Each group was weighted by the inverse of its first eigenvalue before global singular-value decomposition using prince.MFA. No taxonomic or physiological label was used to construct the axes. Amniota comprised Mammalia, Aves, Reptilia and Lepidosauria.

Amniote versus non-amniote and endothermic versus ectothermic partitions were compared by three complementary measures. The silhouette coefficient was calculated from the ten standardised ratios. Logistic-regression accuracy was evaluated by five-fold stratified cross-validation. PERMANOVA used Euclidean distances in MFA space and 4999 label permutations. These statistics quantify, respectively, cluster compactness, out-of-sample classification and the fraction of multivariate variation associated with the grouping. Non-amniote genomes within or near the amniote cluster were identified from Mahalanobis distance to the amniote centroid under the amniote covariance matrix, using the 95th percentile of the corresponding *χ*^2^ distribution as the threshold.

### Isochore analysis

Sixteen assemblies, comprising ten amniotes and six non-amniotes, were partitioned into 100-kb windows without overlap. Windows were retained when at least 50 % of bases were unambiguous; more than 96 % of windows were retained in every assembly. Soft-masked lower-case bases were counted as unambiguous. Each window was assigned a G+C content and a region-specific *ρ*_CG_, then classified into a Bernardi family: L1, *<* 37 %; L2, 37–*<* 41 %; H1, 41–*<* 46 %; H2, 46–*<* 53 %; and H3, *≥* 53 % G+C.

Genome-average G+C, isochore-family coverage and family-specific median *ρ*_CG_ were compared between amniotes and non-amniotes by Mann–Whitney tests across genomes. Monotonic change in *ρ*_CG_ across the ordered families was assessed by Spearman correlation separately for each genome and in the pooled data.

### Invertebrate comparative control

The complete counting and nested-model coupling analysis was repeated on the 481 invertebrate genomes without changing coordinate definitions, predictors or model scoring. Leave-one-order-out cross-validation again used taxonomic order as the held-out unit. The invertebrate results were compared with the vertebrate results as a test of whether strong CpG depletion and AG/CT coupling are general properties of animal genomes.

### Divergence-age summary and compositional topology

For the exploratory divergence-age comparison, median *ρ*_CG_ and sample size were calculated for each of ten vertebrate classes. Stem divergence ages were obtained from TimeTree 5 (24), and their association with class median *ρ*_CG_ was evaluated by Spearman correlation. The class values were not treated as phylogenetically independent.

For the topology analysis, genomes were aggregated by taxonomic order. The 60 orders with at least three genomes and a placement in NCBI Taxonomy were retained. Each order was represented by the mean of its ten ratios after genome-level z-scoring. The reference vertebrate topology was obtained from NCBI Taxonomy (25); orders were placed within their classes and intermediate unary nodes were collapsed. The compositional dendrogram was built by Ward linkage on Euclidean distances among standardised order means. Agreement was quantified by a Mantel test between the cophenetic distance matrices of the two trees with 4999 permutations.

The tanglegram places the two trees face to face and connects identical orders with class- coloured lines. The heatmap displays the ten z-scored ratios in compositional-dendrogram order. The radial reference tree adds concentric rings for class, mean *ρ*_CG_ and z-scored AG/CT.

### Vertebrate–invertebrate machine-learning analysis

The deduplicated vertebrate and invertebrate datasets were combined and represented by the ten strand-symmetric *ρ* coordinates. A random forest of 1000 trees with balanced class weights was evaluated by stratified ten-fold cross-validation. We report pooled out-of-fold ROC and AUC, accuracy, the confusion matrix, Matthews correlation coefficient, impurity importance and held-out permutation importance. A principal component analysis was performed on the ten standardised coordinates. Two reduced models were also evaluated: logistic regression using *ρ*_CG_ alone and a random forest trained on the other nine coordinates after *ρ*_CG_ was removed. Because folds were stratified by class rather than blocked by taxonomic lineage, these metrics quantify discrimination among the sampled genomes and not generalisation to entirely unseen clades.

## Results

### Strand symmetry and base-composition normalisation define the dinucleotide coordinates

We first defined a coordinate system that separates dinucleotide-specific variation from differences in base composition among genomes. This required two distinct operations: testing whether reverse-complement pairs could be combined and normalising each resulting coordinate to its base-composition expectation. For the first operation, we asked whether intrastrand reverse-complement symmetry is sufficiently accurate to reduce the sixteen dinucleotides to ten coordinates. This symmetry is not implied by DNA complementarity alone: complementarity relates a sequence on one strand to its reverse complement on the opposite strand, whereas the present analysis requires the two sequences to have similar frequencies within the same strand. Across the 753 vertebrate nuclear genomes, the frequencies of the two members of each reverse-complement pair were nevertheless almost identical (Pearson *r* = 0.9998; mean absolute frequency difference = 1.4 *×* 10*^−^*^4^). We therefore retained AT, TA, CG and GC individually and averaged the members of the six pairs AA/TT, AC/GT, AG/CT, CA/TG, GA/TC and CC/GG (see Materials and Methods).

For the second operation, we expressed each coordinate as an observed-over-expected ratio, *ρ*. For a dinucleotide *XY*, *ρ_XY_* is its observed frequency divided by *f_X_f_Y_*, the frequency expected under independent occurrence of the two bases while retaining that genome’s base composition. This ratio answers the biologically relevant question: does *XY* occur more or less often than expected in a genome with the same proportions of *X* and *Y*A value of one indicates no enrichment or depletion relative to base composition; values below and above one indicate depletion and enrichment, respectively. Unlike raw dinucleotide frequency, *ρ* therefore permits direct comparisons among genomes with different A, T, G and C contents. Pairing reduces redundant strand orientation, whereas the *ρ* normalisation provides the quantity interpreted throughout the analysis. The CpG coordinate is denoted *ρ*_CG_ (see Materials and Methods).

Vertebrate genomes were strongly depleted of CpG (Figure 1). Across genomes, CpG had a mean observed frequency of 1.53 %, compared with 4.41 % expected from base composition. The mean genome-level *ρ*_CG_ was therefore 0.345, indicating that vertebrate genomes retain, on average, approximately one third of the CpG dinucleotides expected under the base-composition model.

**Figure 1:**
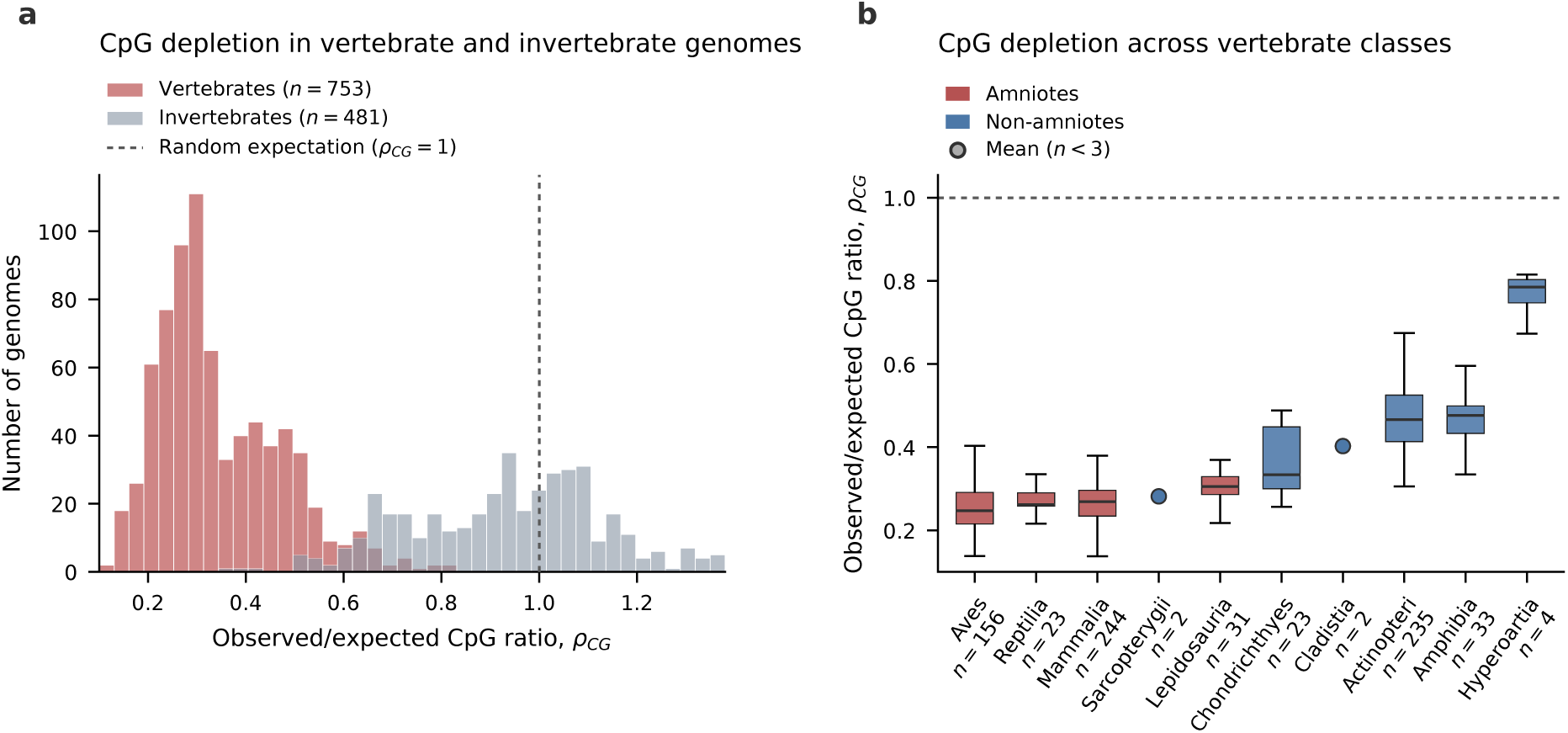
Genome-wide CpG depletion in vertebrates. The quantity plotted is the CpG observed-over-expected ratio *ρ*_CG_, the number of CG dinucleotides present in a genome divided by the number expected if its bases were arranged at random for the same base composition; a value of one means no depletion, and values below one mean CpG is rarer than chance. **(A)** Distribution of *ρ*_CG_ across 753 vertebrate and 481 invertebrate genomes; vertebrates are strongly depleted (mean 0.345), whereas invertebrates centre near the no-bias value of one (vertical dotted line). **(B)** The same ratio by vertebrate class, ordered by median and coloured by clade, amniote against non-amniote, showing the gradient from heavily eroded amniotes to nearly unbiased jawless fishes (the no-bias value of one is the horizontal dotted line). Classes with fewer than three genomes are drawn as a single mean point.

Because related genomes are not statistically independent (18), uncertainty was estimated by resampling taxonomic orders rather than individual assemblies. In each of 2000 clade-block bootstrap replicates, complete orders and all their constituent genomes were sampled with replacement (see Materials and Methods). The resulting 95 % confidence interval for mean *ρ*_CG_ was 0.31–0.39 (Supplementary Table S1 and Supplementary Figure S1). This interval is wider, and more appropriate for the phylogenetically structured dataset, than an ordinary genome-level bootstrap interval.

The ten-coordinate representation also has a mechanistic basis. If the complete context-dependent mutation process is identical on both strands, the mutation–replication dynamics are invariant under reverse complementation; at equilibrium, a sequence and its reverse complement consequently have the same expected frequency on either strand. A two-strand simulation initiated from a deliberately asymmetric sequence converged to reverse-complement parity under symmetric rates, whereas the measured strand bias of human mitochondrial DNA prevented convergence. The complete simulation, including its assumptions and controls, is reported in Supplementary Methods and Results and Supplementary Figures S2 and S3. This analysis justifies the ten-coordinate representation for the nuclear genomes considered here while defining the condition under which it would fail.

### AG/CT is the strongest between-genome signature of CpG loss

The next question was which coordinate most faithfully records differences in CpG depletion among vertebrate genomes. We first tested the direct products of methyl-CpG deamination. Under strand symmetry, a fixed CpG-to-TpG/CpA event contributes to the two members of the CA/TG coordinate in opposite orientations. Because this coordinate is their mean, its expected excess is one half of the CpG deficit (see Materials and Methods). Across the complete vertebrate set, the observed ratio of mean TpG/CpA excess to mean CpG deficit was 0.49 (clade-block bootstrap 95 % confidence interval 0.45–0.55), consistent with the expected value of 0.5 (Supplementary Table S1 and Supplementary Figure S1). Thus, the dataset-wide chemical balance is recovered.

The same coordinate was a poor marker of variation among individual genomes. CA/TG correlated only weakly with *ρ*_CG_ (Pearson *r* = +0.27), indicating that a genome with greater CpG depletion does not necessarily have a proportionally greater CA/TG ratio. TpG and CpA are abundant and have multiple sequence-context sources; the CpG-derived component can therefore satisfy the mean mass balance while remaining difficult to resolve against the genome-specific background.

We consequently tested all other strand-symmetric coordinates. AG/CT showed the strongest relationship: genomes with lower *ρ*_CG_, and hence greater CpG depletion, had higher AG/CT (Pearson *r* = *−*0.92; Figure 2). AC/GT also changed with *ρ*_CG_, but in the same direction (Pearson *r* = +0.71). Neither AG/CT nor AC/GT is an immediate product of CpG deamination, suggesting that CpG loss propagates beyond TpG/CpA through context-dependent mutation.

**Figure 2:**
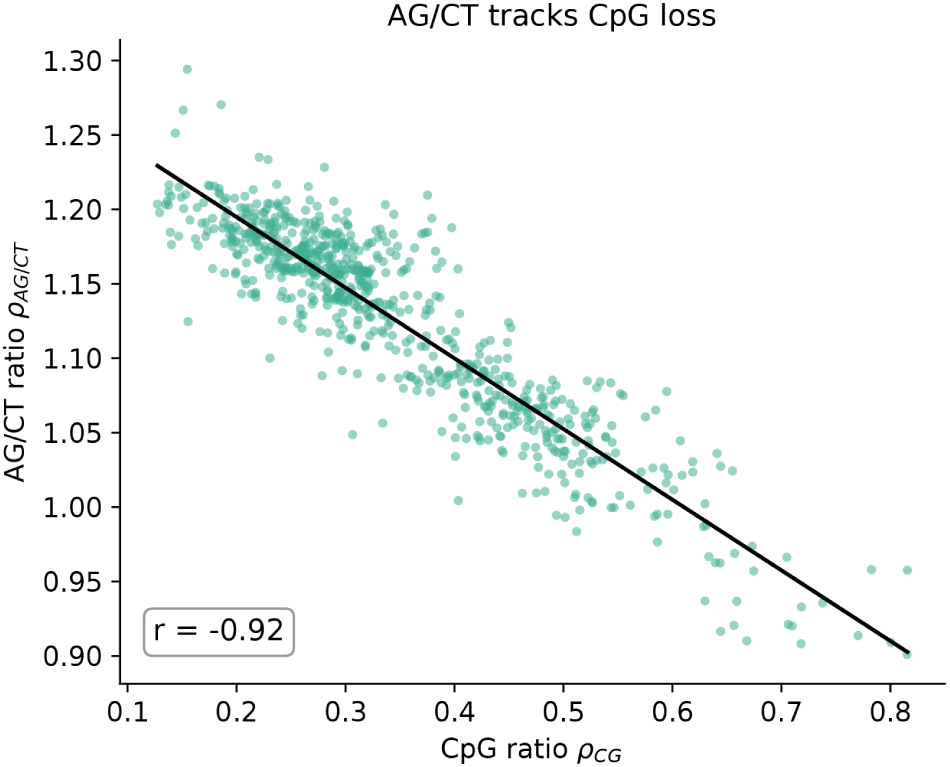
AG/CT is the strongest between-genome signature of CpG depletion. Each point represents one of the 753 vertebrate genomes. Lower *ρ*_CG_ denotes stronger CpG depletion; AG/CT increases as *ρ*_CG_ decreases (Pearson *r* = *−*0.92).

To distinguish CpG-specific coupling from a shared dependence on base composition, we compared two nested models for each non-CG coordinate. The baseline model used only the abundance expected from the component bases; the augmented model also included the observed CpG frequency. Their difference, Δ*R*^2^, measures the additional between-genome variance explained by CpG after base composition has been taken into account (see Materials and Methods). AG/CT had the largest gain (Δ*R*^2^ = 0.74), which increased to 0.82 under leave-one-order-out cross-validation. AC/GT ranked second (Δ*R*^2^ = 0.50), whereas the direct-product coordinate CA/TG gained only 0.05 (Table 1; complete statistics in Supplementary Table S2 and Supplementary Figure S4). Dataset-wide mass balance and between-genome association thus answer different questions: CA/TG recovers the average product of CpG loss, whereas AG/CT best distinguishes genomes with different levels of CpG depletion.

**Table 1:** Coupling of each strand-symmetric coordinate to CpG across 753 vertebrate genomes. “Base only” is the variance explained by base composition; “Base + CpG” is the variance explained after observed CpG frequency is added. Δ*R*^2^ is the gain attributable to CpG, irrespective of the direction of association. The final column reports the gain under leave-one-order-out cross-validation.

| Coordinate | Base only | Base + CpG | $\Delta R^2$ | $\Delta R^2_{cv}$ |
| --- | --- | --- | --- | --- |
| AG/CT | 0.15 | 0.89 | 0.74 | 0.82 |
| AC/GT | 0.00 | 0.50 | 0.50 | 0.53 |
| CC/GG | 0.61 | 0.78 | 0.17 | 0.18 |
| GA/TC | 0.07 | 0.19 | 0.11 | 0.12 |
| CA/TG | 0.09 | 0.14 | 0.05 | 0.03 |
| AT | 0.73 | 0.77 | 0.04 | 0.04 |
| TA | 0.61 | 0.62 | 0.02 | 0.00 |
| AA/TT | 0.78 | 0.79 | 0.01 | 0.00 |
| GC | 0.55 | 0.55 | 0.00 | -0.02 |

Finally, we tested whether the AG/CT association could be explained by compositional closure or taxonomic non-independence. The correlation with *ρ*_CG_ remained *−*0.92 (clade-block bootstrap 95 % confidence interval *−*0.94–*−*0.87), and was *−*0.87 after a centred-log-ratio transformation of the complete sixteen-part dinucleotide composition. It also remained significant in a taxonomy- structured generalised least-squares model and after aggregation by order or class (see Materials and Methods). By contrast, the AC/GT association became negligible after taxonomic relatedness was included, indicating that it is largely a between-clade pattern. AG/CT is therefore the most robust second-order signature of CpG loss (Supplementary Table S3 and Supplementary Figure S5).

### Context-dependent mutation explains the AG/CT signal

The correlation analysis established that AG/CT tracks CpG depletion, but not why. We therefore asked whether the AG/CT increase can be generated by CpG loss propagating through a measured context-dependent mutation network. The test was deliberately predictive: if both signals arise from the same process, a model calibrated to CpG alone should recover AG/CT without an AG/CT-specific parameter.

We evolved random sequences using Trek estimates of human germline substitution rates (17). These rates specify the three possible substitutions of a central base within each seven-nucleotide context, comprising the central base and its three flanking bases on either side (see Materials and Methods). Before adding a CpG-specific term, we validated the implementation by starting from sequences with 20 % or 60 % G+C. Both converged to the published equilibrium of 38.2 % G+C and were continued to approximately five substitutions per site (Supplementary Figure S6; see Materials and Methods). This convergence shows that the numerical engine reproduces the equilibrium of the input rates independently of starting composition.

The Trek core rates intentionally exclude the elevated mutability of methylated CpG. Under these rates alone, the simulated sequence remained essentially undepleted for CpG (*ρ*_CG_ = 1.052) and AG/CT remained close to one (*ρ*_AG_*_/_*_CT_ = 1.028). We then introduced a single multiplier, *m*, for C-to-T substitution at CpG sites and the reverse-complementary G-to-A event. The multiplier was calibrated only to the mammalian median *ρ*_CG_ = 0.269, giving *m* = 7.91 and a simulated *ρ*_CG_ = 0.271 (see Materials and Methods).

AG/CT provided the independent test. No parameter was fitted to this coordinate. Once the calibrated CpG term was included, simulated AG/CT increased from 1.028 to 1.138, close to the observed mammalian median of 1.160 (Figure 3A). The change arises because a CpG-to- TpG mutation alters the sequence contexts, and therefore the subsequent substitution rates, of neighbouring positions. Repeated context changes transmit the initial CpG perturbation through the mutation network. The model therefore explains AG/CT as a second-order consequence of methylation-associated CpG loss rather than as a direct deamination product.

**Figure 3:**
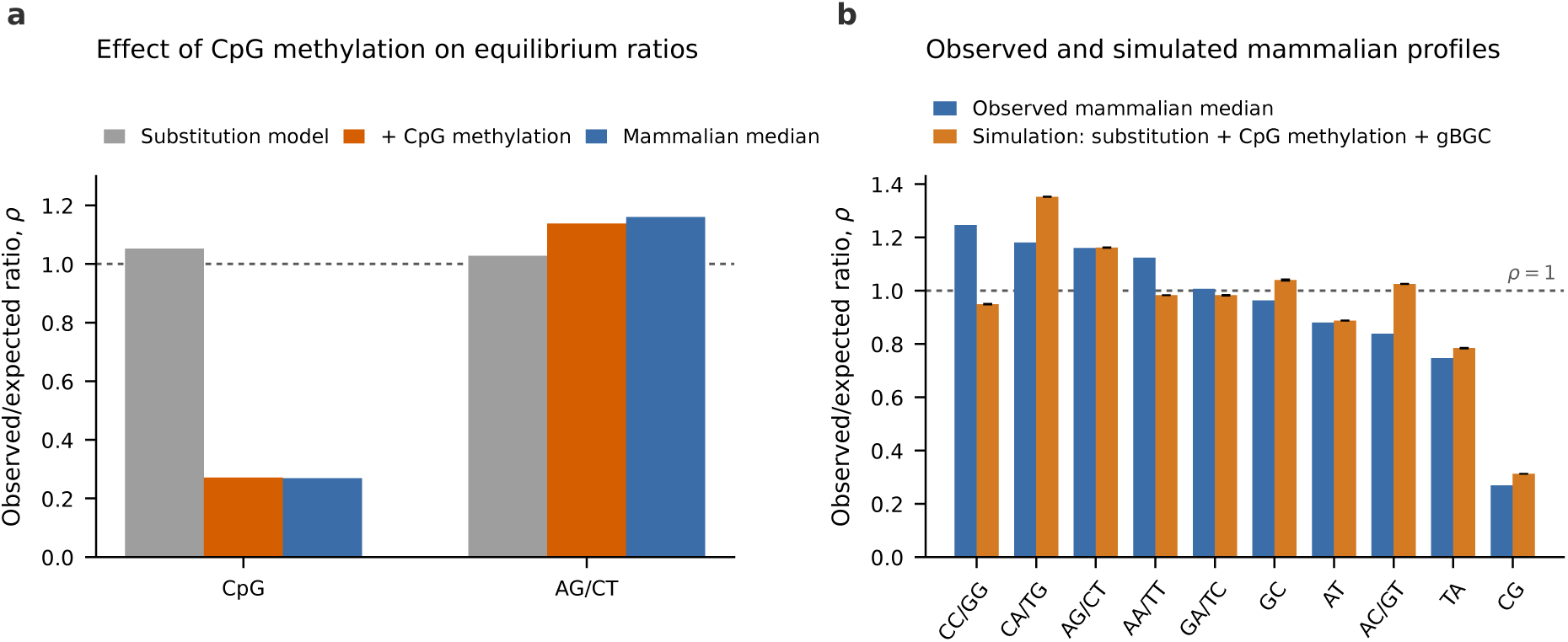
A CpG-specific methylation term reproduces AG/CT enrichment. **(A)** Equilibrium *ρ* values for CpG and AG/CT under the Trek core rates (grey), after addition of the calibrated CpG-mutability multiplier *m* = 7.91 (orange), and in mammalian genomes (blue). Only CpG was used for calibration; AG/CT is an independent model prediction. **(B)** Mammalian medians and simulated equilibrium values for all ten strand-symmetric coordinates under the complete model, which includes context-dependent substitution, CpG methylation and the GC-biased-gene-conversion approximation. The dashed line denotes *ρ* = 1.

The mutation-plus-methylation model equilibrated at approximately 38.2 % G+C, below the 41.0 % of the human assembly. To represent GC-biased gene conversion (gBGC), which favours G and C alleles during heteroduplex repair (19), we added a second factor, *b*, to every substitution producing G or C. Calibrating only to human G+C content gave *b* = 1.50 (see Materials and Methods). This term altered base composition and shifted some individual *ρ* values, but preserved the principal CpG–AG/CT pattern (Figure 3B).

Across the ten coordinates, frequencies from the complete model and the T2T-CHM13 human reference genome correlated at *r* = 0.94, with a mean absolute error of 0.53 percentage points (Figure 4A). On the *ρ* scale, AG/CT was recovered to within 0.001 of the observed value (Figure 4B and Supplementary Table S4). The largest residuals were the underprediction of AA/TT (*−*0.14) and CC/GG (*−*0.30). A substitution-only model cannot lengthen homopolymer runs, whereas replication slippage and short insertion/deletion events can do so at rates far above the point-mutation rate at these loci (20, 21). Because slippage is not required to test the CpG-to-AG/CT mechanism, no additional parameter was fitted.

**Figure 4:**
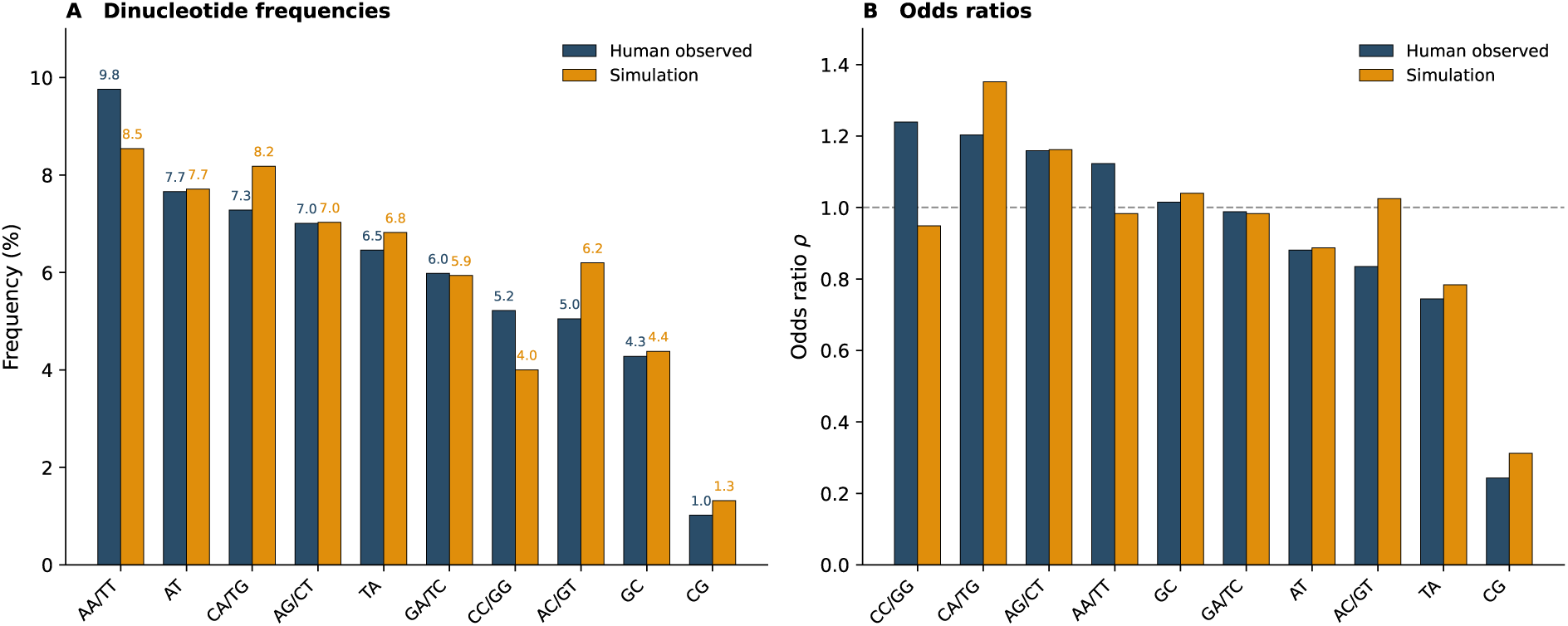
Observed and simulated human dinucleotide profiles. **(A)** Strand-symmetric dinucleotide frequencies in the human genome (blue) and at equilibrium under the complete model (orange), combining seven-mer substitution rates, CpG methylation (*m* = 7.91) and the gBGC approximation (*b* = 1.50). Pearson *r* = 0.94; mean absolute error = 0.53 percentage points. **(B)** The same comparison on the *ρ* scale. AG/CT is recovered to within 0.001; AA/TT and CC/GG are underpredicted because replication slippage is not included.

### CpG depletion is concentrated within transposable elements

We next asked where the CpG-depletion signal is located within vertebrate genomes. RepeatMasker annotations were available and sufficiently complete for pooled testing in 11 genomes (see Materials and Methods). Median *ρ*_CG_ was lowest inside transposable elements (0.19) and increased with distance from the nearest element, reaching 0.39 beyond 5 kb (Figure 5). Thus, the *severity of CpG depletion decreases* with distance: the observed-over-expected ratio itself increases. This direction is important because low *ρ*_CG_, not high *ρ*_CG_, denotes stronger depletion.

**Figure 5:**
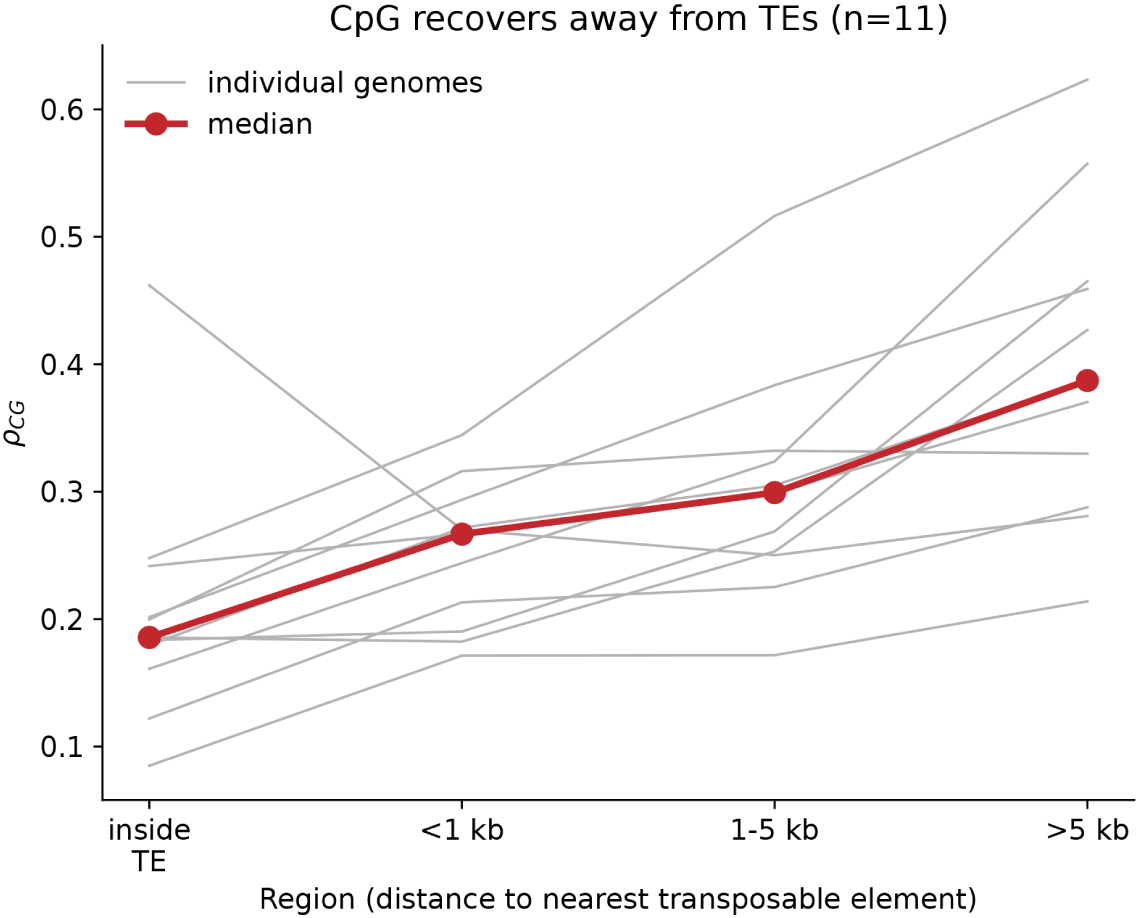
CpG depletion is strongest inside transposable elements and weakens with distance. The CpG observed-over-expected ratio is lowest within elements and increases with distance from the nearest element. Grey lines represent individual genomes; the red line is their median.

Inside-element *ρ*_CG_ was lower than the value beyond 5 kb in 10 of 11 genomes (two-sided exact Wilcoxon signed-rank *p* = 0.0098). Across genomes, transposable-element content and *ρ*_CG_ had a partial correlation of *−*0.29 after AG/CT was held constant, so genomes with more annotated element sequence tended to be more CpG-depleted. Together, the regional and between-genome results localise the strongest CpG loss to transposable elements, consistent with the long-term mutational effect of their methylation-mediated silencing. Per-genome values, region definitions and assembly provenance are provided in Supplementary Tables S5 and S6.

### The compositional division follows Amniota more closely than endothermy

We then asked whether the principal compositional structure among vertebrates is associated more strongly with common ancestry or with thermophysiology. Multiple factor analysis (MFA) was performed on two standardised variable groups: base composition and the ten dinucleotide ratios. Taxonomic and physiological labels were not used to construct the axes (see Materials and Methods). The first two dimensions accounted for 30.4 % and 28.9 % of the total variation, respectively (Figure 6 and Supplementary Table S7). We compared two external partitions of this same unsupervised space: amniotes versus non-amniotes and endotherms versus ectotherms.

**Figure 6:**
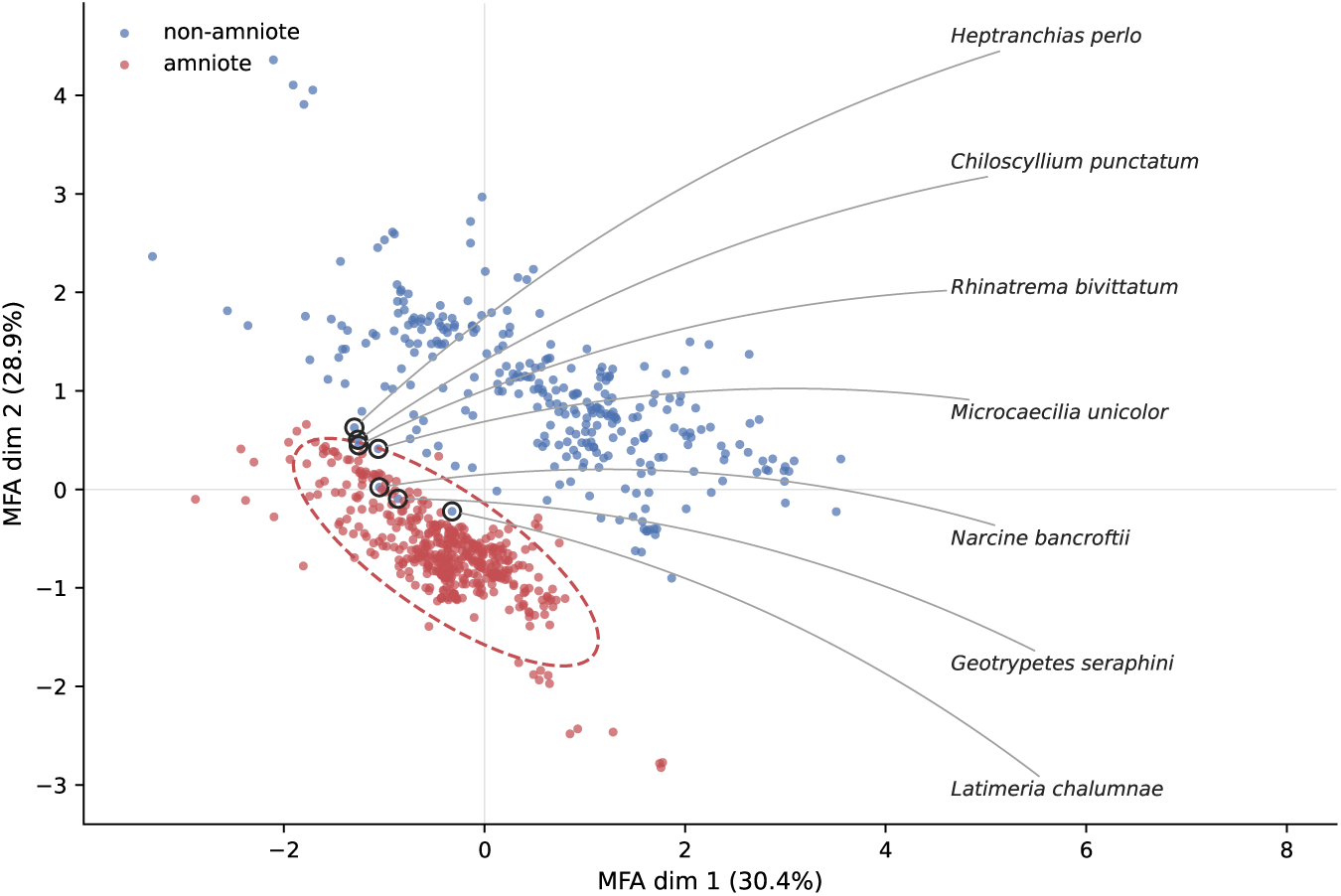
Multiple factor analysis of vertebrate base and dinucleotide composition. The axes were constructed from standardised base frequencies and the ten dinucleotide ratios, without taxonomic or physiological labels. Points are coloured by clade; the dashed ellipse is the 95 % amniote confidence ellipse. Labels identify the seven non-amniote genomes closest to the amniote centroid by Mahalanobis distance. Dimensions 1 and 2 account for 30.4 % and 28.9 % of the variation, respectively.

All three separation measures favoured the amniote partition. The silhouette coefficient based on the ten standardised ratios was 0.371 for amniote status and 0.284 for thermophysiology. Five-fold cross-validated logistic-regression accuracy was 0.993 and 0.954, respectively. PERMANOVA on the MFA coordinates attributed *R*^2^ = 0.260 (pseudo-*F* = 263) to amniote status, compared with *R*^2^ = 0.211 (pseudo-*F* = 201) for thermophysiology (see Materials and Methods).

Ectothermic amniotes separate these alternatives most clearly. Their median *ρ*_CG_ was 0.291, close to that of endothermic amniotes (0.264) and markedly below that of ectothermic fishes and amphibians (approximately 0.44). If endothermy were the main determinant, ectothermic amniotes would be expected to group with other ectotherms. Instead, they grouped with mammals and birds. The compositional pattern is therefore more consistent with an amniote-associated state than with endothermy, which originated independently in mammals and birds.

The separation was not absolute. On the basis of Mahalanobis distance from the amniote centroid, seven non-amniote genomes fell within or near the amniote cluster. Three, *Geotrypetes seraphini*, *Narcine bancroftii* and *Latimeria chalumnae*, fell within it, whereas *Rhinatrema bivittatum*, *Microcaecilia unicolor*, *Chiloscyllium punctatum* and *Heptranchias perlo* lay near its boundary. These exceptions show that the compositional transition is graded rather than a strict taxonomic boundary; they do not alter the stronger overall fit of the amniote partition.

### Regional GC organisation reveals an amniote signal hidden by genome averages

We next asked whether the amniote-associated pattern is visible in regional genome composition even though mean G+C content is nearly identical in amniotes and non-amniotes (42.3 % versus 42.0 %; Mann–Whitney *p* = 0.87). Sixteen assemblies were divided into non-overlapping 100-kb windows and each window was assigned to a Bernardi isochore family, from GC-poor L1 to GC-rich H3 (see Materials and Methods). H3 windows covered 3.9 % of the ten amniote genomes and 1.3 % of the six non-amniote genomes, a 3.0-fold difference (Mann–Whitney *p* = 0.022; Figure 7A).

**Figure 7:**
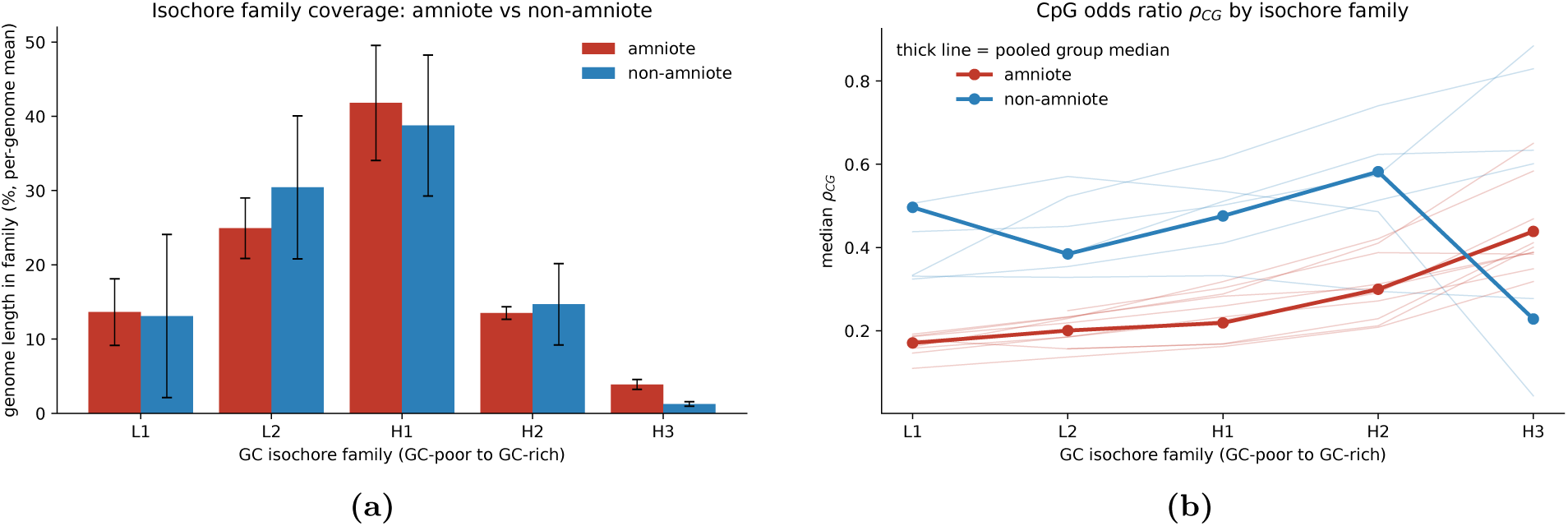
Regional GC organisation distinguishes amniotes despite similar genome-average G+C. **(A)** Fraction of sequence assigned to each isochore family in amniotes and non-amniotes. GC-rich H3 sequence is 3.0 times more extensive in amniotes. **(B)** Median *ρ*_CG_ by isochore family. In amniotes, *ρ*_CG_ rises from 0.171 in L1 to 0.439 in H3, indicating progressively weaker CpG depletion.

CpG depletion also differed among these compartments. In amniotes, median *ρ*_CG_ increased from 0.171 in L1 windows to 0.439 in H3 windows (Figure 7B and Supplementary Table S8). Because higher *ρ*_CG_ means weaker depletion, CpG loss was greatest in GC-poor regions and progressively weaker in GC-richer regions. This monotonic pattern was observed in all ten amniote genomes.

An increase in *ρ*_CG_ from GC-poor to GC-rich windows was also present in four of the six non-amniote genomes and is therefore not exclusive to Amniota. The groups differed in the magnitude and consistency of the gradient and in the extent of the H3 compartment. Amniotes had substantially lower *ρ*_CG_ in L1, displayed the gradient in every genome examined and contained 3.0 times more H3 sequence. Their distinguishing feature is therefore regional genome organisation rather than higher mean G+C: a strongly CpG-depleted GC-poor compartment coexists with a larger GC-rich compartment in which CpG depletion is partially reduced.

The regional comparison includes only ten amniote and six non-amniote genomes and should be interpreted accordingly. Its possible connection to higher-order genome organisation is also hypothesis-generating rather than tested here. GC-rich isochores have been associated with RNA processing through nuclear speckles (22), and related sequence features have been linked to three-dimensional chromosome organisation (23).

### The CpG–AG/CT axis is attenuated in invertebrates

We used the 481 invertebrate genomes to ask whether the vertebrate CpG–AG/CT relationship is a general property of animal genomes. The same counting and CpG-coupling pipeline was applied without alteration (see Materials and Methods). Mean invertebrate *ρ*_CG_ was 1.02, close to the no-depletion value of one. AG/CT coupling was also much weaker than in vertebrates (Δ*R*^2^ = 0.23 versus 0.74), and the largest invertebrate gain occurred for CA/TG (Δ*R*^2^ = 0.42; Supplementary Figure S7 and Supplementary Table S9). The vertebrate signatures were therefore attenuated rather than universally absent. Their weaker and more heterogeneous expression is consistent with the diversity of invertebrate methylation systems among phyla (12–14).

### Dinucleotide composition retains phylogenetic structure

We next asked how closely the ten-coordinate profile follows vertebrate ancestry. As an exploratory summary, class median *ρ*_CG_ was compared with stem divergence age from TimeTree 5 (24). The association was positive for *ρ*_CG_, meaning that classes with older stem ages tended to show *weaker*, not stronger, CpG depletion (Supplementary Figure S8 and Supplementary Table S10). This comparison contains only ten non-independent class values, is sensitive to Hyperoartia and should not be interpreted as a temporal rate of CpG loss.

For a direct topology comparison, we averaged the ten standardised ratios within the 60 taxonomic orders represented by at least three genomes, constructed an unsupervised Ward dendrogram and compared it with the NCBI Taxonomy reference tree (25) (see Materials and Methods). Cophenetic distances from the two trees were correlated (Mantel *r* = 0.735, *p <* 0.001), showing substantial but incomplete phylogenetic structure (Supplementary Figure S9). Related orders often had similar profiles, but amniote orders were also drawn together by their shared CpG depletion and AG/CT increase rather than solely by their positions in the reference topology.

The heatmap of z-scored coordinates identifies the compositional features underlying these clusters (Supplementary Figure S10). A radial representation of the reference topology reaches the same conclusion: the rings for mean *ρ*_CG_ and z-scored AG/CT follow the amniote lineages while also varying across the broader tree (Supplementary Figure S11). Dinucleotide composition therefore contains phylogenetic information, but a shared CpG-depletion regime can increase similarity across branches and prevents composition from serving as a direct substitute for the reference topology.

### CpG and AG/CT provide the principal vertebrate–invertebrate classification signal: a Machine Learning analysis

Finally, we asked whether the vertebrate compositional regime is distributed across many independent coordinates or is primarily encoded by the CpG–AG/CT axis. A class-balanced random forest using all ten ratios was evaluated by stratified ten-fold cross-validation (see Materials and Methods). Pooled out-of-fold predictions achieved an AUC of 0.999, accuracy of 0.983 and Matthews correlation coefficient of 0.964 (Supplementary Figure S12 and Supplementary Table S11).

The confusion matrix contained 14 vertebrate false negatives and 7 invertebrate false positives (Supplementary Figure S13). False-negative vertebrates included pipefishes, seahorses and lampreys, which have comparatively weak CpG depletion. False-positive invertebrates were unusually CpG-poor. The errors therefore occurred mainly in genomes whose CpG profiles resembled the opposite class, rather than being distributed randomly.

We evaluated the importance of individual coordinates within this classification task in two complementary ways. Within the ten-feature model, both impurity-based and held-out permutation importance ranked *ρ*_CG_ first and AG/CT second (Supplementary Figure S14). In a separate single-coordinate test, logistic regression using *ρ*_CG_ alone reached an AUC of 0.992, close to the AUC of 0.999 obtained with all ten coordinates (Supplementary Table S11). These results identify *ρ*_CG_ as the strongest individual discriminator of the sampled vertebrate and invertebrate genomes.

This importance is not unique, because the coordinates carry correlated information. A forest trained on the other nine ratios after *ρ*_CG_ was removed retained an AUC of 0.999. A principal component analysis provided an unsupervised view of the same structure: PC1 explained 41.6 % of the variance, PC2 explained 20.7 %, and CpG and AG/CT had opposing loadings on the principal axis separating the two groups (Supplementary Figure S15). We therefore use “dominant” in a specific empirical sense: CpG is the leading single marker of a coordinated CpG–AG/CT compositional axis in these data. The results do not imply that CpG uniquely determines genome composition or independently explains every dinucleotide coordinate.

## Discussion

The main result is that vertebrate dinucleotide composition contains a coordinated axis associated with CpG loss, and that the strongest secondary marker of this axis can be explained mechanistically. Previous models showed that CpG-to-TpG/CpA mutation can alter multiple dinucleotides and genomic G+C (26). The present analysis extends that framework in two ways: it identifies AG/CT as the most robust between-genome signature after explicit compositional and taxonomic controls, and it shows that the observed AG/CT value emerges from measured seven-mer substitution rates when a CpG-specific methylation term is added.

No single result proves that CpG is a universal determinant of genome composition. The description of a dominant CpG-associated axis is instead supported by three convergent analyses. First, adding observed CpG frequency to the base-composition model increased the explained variance of AG/CT by 0.74, rising to 0.82 under leave-one-order-out validation (Table 1). Second, the methylation multiplier was calibrated only to *ρ*_CG_; without fitting AG/CT, it raised the simulated AG/CT value from 1.028 to 1.138, close to the mammalian median of 1.160 (Figure 3). After the separate G+C calibration, the complete human model recovered AG/CT to within 0.001 of the observed value (Figure 4). Third, *ρ*_CG_ ranked first in the vertebrate–invertebrate feature-importance analysis and achieved an AUC of 0.992 as the only predictor, while CpG and AG/CT loaded in opposite directions on the leading principal component. Together, these results support CpG as the leading individual coordinate of the dinucleotide variation examined here; they do not establish CpG as a general model of every aspect of genome composition or structure. The reduction from sixteen dinucleotides to ten is supported both empirically and mathematically. Complementarity alone relates opposite strands; it does not require a sequence and its reverse complement to be equally abundant within one strand. If the complete context-dependent mutation process is the same on both strands, however, the mutation–replication dynamics are invariant under reverse complementation. Their equilibrium distribution must therefore satisfy reverse-complement parity. The two-strand simulation confirms this convergence from an asymmetric initial condition and shows that measured mitochondrial strand bias prevents it. This is consistent with general no-strand-bias models (5, 27, 28); inversion and duplication may provide an additional route to the same symmetry (8). The ten-coordinate representation is consequently appropriate for the nuclear genomes analysed here, but should not be applied without verification to persistently strand-biased genomes.

The distinction between TpG/CpA and AG/CT is central. TpG/CpA satisfies the expected mean mass balance across the vertebrate dataset, supporting its role as the direct product of CpG loss. However, it varies only weakly with *ρ*_CG_ among genomes. The direct-product background is large and has multiple sources, so a correct global balance need not provide a sensitive genome-level marker. AG/CT answers a different question: it captures how the broader dinucleotide network differs among genomes with different amounts of CpG depletion. Its association remains strong after adjustment for base composition, compositional closure and taxonomic structure; the corresponding AC/GT signal is largely between clades.

The forward model provides the mechanistic link. Core human germline rates generate neither substantial CpG depletion nor AG/CT enrichment. A methylated-CpG multiplier calibrated only to *ρ*_CG_ raises simulated AG/CT from 1.028 to 1.138, without fitting AG/CT itself. After the separate gBGC term is calibrated to human G+C content, the complete model recovers AG/CT to within 0.001 of the observed human value. AG/CT is therefore transmitted through changes in neighbouring sequence context rather than produced directly by CpG chemistry, consistent with the capacity of context-dependent mutation networks to generate large-scale sequence regularities (5, 8). The result is strongest as a mechanistic demonstration for the human rate system. Seven-mer rates of comparable resolution are not available for the other species, so extension to all vertebrates is supported by the comparative association, not by species-specific forward models.

The model is intentionally incomplete. Its gBGC term is a simplified approximation calibrated to human G+C content, and the model reaches equilibrium rather than reconstructing a dated evolutionary trajectory. It also omits replication slippage. The resulting underprediction of AA/TT and CC/GG is therefore informative rather than unexpected: substitution alone cannot extend homopolymer tracts, whereas slippage can occur at rates far exceeding point mutation (20, 21). Adding a fitted slippage parameter would improve those coordinates but would not test the proposed CpG-to-AG/CT mechanism.

The spatial analysis gives the axis a genomic location. *ρ*_CG_ is lowest within transposable elements and rises with distance, meaning that CpG depletion is strongest inside elements and weakens away from them. This pattern is consistent with methylation-mediated element silencing followed by long-term CpG-to-TpG/CpA mutation, and agrees with evidence connecting methylation and transposable-element content across animals (13, 14). The analysis does not measure present-day methylation directly, and the pooled test is restricted to eleven genomes with sufficient repeat annotation; it should therefore be interpreted as a compositional footprint consistent with that mechanism rather than a direct methylation assay.

Across vertebrates, the same axis follows Amniota more closely than endothermy. Ectothermic amniotes group with mammals and birds rather than with ectothermic fishes and amphibians, and every separation metric favours the amniote partition. Because endothermy originated independently in mammals and birds whereas Amniota has a single origin, the more parsimonious interpretation is an amniote-associated germline compositional regime retained across descendant lineages. The graded boundary and the non-amniote genomes near the amniote cluster show that this is not a deterministic taxonomic classifier.

Genome-average G+C does not explain this division: amniotes and non-amniotes have almost identical mean G+C, but differ in its regional distribution. Amniotes contain a larger GC-rich H3 compartment, and their *ρ*_CG_ rises from GC-poor to GC-rich isochores. gBGC is a plausible contributor to this regional organisation, but the simple simulation term does not establish that mechanism. Likewise, connections to nuclear speckles and three-dimensional organisation (22, 23) remain hypotheses because those data were not analysed here.

Finally, the partial correspondence between the compositional dendrogram and the reference topology shows that the ten ratios retain phylogenetic information while also reflecting a shared methylation-associated state. The classifier reaches the same conclusion from a different direction: *ρ*_CG_ and AG/CT dominate vertebrate–invertebrate separation, but either *ρ*_CG_ alone or the remaining correlated coordinates retains most predictive information. The weak mean CpG depletion and reduced AG/CT coupling of the heterogeneous invertebrate set further delimit the result. The axis described here is a characteristic of the pervasive vertebrate regime, not a universal rule of animal genome composition.

## Conclusions

Across 753 vertebrate genomes, CpG depletion defines a coordinated axis of dinucleotide variation. Its most informative between-genome signature is not the direct product TpG/CpA, which recovers the expected mean mass balance, but AG/CT. Forward evolution under seven-mer human germline substitution rates shows why: a CpG-mutability term fitted only to the mammalian CpG ratio also predicts mammalian AG/CT. The AG/CT increase is therefore a mechanistically predictable, second-order output of the context-dependent mutation network.

The same signal is spatially and evolutionarily structured. CpG depletion is strongest within transposable elements and weakens with distance from them. Across lineages, dinucleotide composition follows Amniota more closely than endothermy and is associated with a larger GC-rich isochore compartment that is invisible in genome-average G+C. Composition retains substantial phylogenetic structure, but shared CpG depletion also draws lineages together across the reference tree.

Finally, the CpG–AG/CT axis is markedly weaker in invertebrates and supplies most of the information used to distinguish the two animal groups. Vertebrate dinucleotide frequencies should therefore not be interpreted as ten independent properties: after base composition is accounted for, they largely record a shared, methylation-associated evolutionary process.

## Data availability

Analysis code is available at https://github.com/tacc-code-tacclab/CGtrek. Versioned NCBI RefSeq accessions and annotation releases for the transposable-element analysis are listed in Supplementary Table S6; accession-level provenance for the isochore analysis is provided in the repository.

## Supplementary Information

*Guide to this document.* This section contains the extended Chargaff-parity analysis and the quantitative material supporting the main text. Numerical results were generated from the two RefSeq count tables (753 vertebrate and 481 invertebrate genomes), the assemblies and RepeatMasker annotations identified in Supplementary Table S6, and the Trek rate tables by the deterministic scripts in the analysis repository.

The quantitative material comprises eleven tables and fifteen figures, numbered in the order in which they are cited from the main and supplementary text.

All genome-derived results depend on the exact assembly and repeat annotation used. To make them reproducible, Supplementary Table S6 lists the versioned NCBI RefSeq accession, the assembly name and the NCBI annotation release, with its date, for every RepeatMasker-annotated genome; the sixteen isochore genomes are recorded in the same form in the repository. The FASTA (<accession>_genomic.fna.gz) and the RepeatMasker output (<accession>_rm.out.gz) were retrieved from the NCBI RefSeq FTP tree for exactly these accessions, and the transposable- element distance bins are fixed as stated in the caption of Supplementary Table S5, so that every per-genome value, including the beyond-5 kb column, is reproducible.

## Supplementary analysis of reverse-complement parity

### Rationale and model

The main analysis combines each dinucleotide with its reverse complement. We therefore tested the condition under which this intrastrand parity follows from the mutation process. Double- stranded complementarity alone guarantees that a sequence on one strand is represented by its reverse complement on the other; it does not, by itself, impose equal frequencies within either strand. If the complete context-dependent mutation process is identical on both strands, however, the mutation–replication dynamics are invariant under reverse complementation. At a unique equilibrium, this invariance requires each sequence and its reverse complement to have the same expected frequency on either strand (5, 27, 28).

The CpG-to-TpG/CpA event illustrates the distinction (Supplementary Figure S2). Deamination changes a methylated cytosine on only one strand and creates a mismatch. If the event is fixed through replication, the daughter molecule carrying the mutation contains TpG in one 5*^′^*–3*^′^* orientation and CpA in the reverse-complement orientation. These are not two simultaneous chemical events. Equal accumulation of TpG and CpA within a single strand emerges across sites and generations when equivalent mutation opportunities occur in both orientations.

To test convergence explicitly, the forward-evolution engine was extended to represent the two DNA strands as separate sequences. A substitution altered only the strand on which it occurred, so the resulting mismatch remained explicit until replication. In every generation, context-dependent rates were read in each strand’s own 5*^′^*–3*^′^* orientation and either strand could mutate. Each parental strand was then copied to form a daughter duplex, and one of the two daughters was retained at random. Only one strand of the retained daughter was counted, matching the convention used for genome assemblies.

The initial sequence was drawn from a first-order Markov chain chosen to violate parity in all six reverse-complement dinucleotide pairs and to contain sufficient CpG for its loss to be observed. Two runs began from this same sequence. In the symmetric regime, the Trek rates and calibrated CpG term were applied identically to both strands. In the strand-biased regime, one strand received the two relative substitution factors measured at fourfold-degenerate sites in human mitochondrial DNA: nine-fold for G-to-A and 1.8-fold for T-to-C on the light strand relative to the heavy strand (30). In the stored orientation of the simulated strand, these corresponded to the reverse-complement C-to-T and A-to-G rates; all other rates were unchanged.

Departure from parity was measured as the mean absolute frequency difference between the members of the six reverse-complement pairs, expressed in percentage points. Because the value fluctuates at equilibrium, it was averaged over the final fifth of each run rather than taken from a single endpoint. The residual variation expected from a finite sequence defines the sampling floor.

### Symmetric mutation produces parity whereas strand bias prevents it

Under identical mutation processes on the two strands, all six reverse-complement pairs converged from the deliberately asymmetric starting state, and their mean difference reached the finite- sequence sampling floor (Supplementary Figure S3A,C). No direct conversion between the members of a pair was required. The result is the numerical convergence predicted by reverse-complement invariance of the mutation–replication process.

When the measured mitochondrial strand biases were introduced with every other setting unchanged, the reverse-complement pairs did not converge to the sampling floor (Supplementary Figure S3B,C). The imposed asymmetry also propagated to contexts that were not directly assigned a biased rate. Thus, strand-symmetric mutation is sufficient to generate dinucleotide parity, whereas persistent strand-specific mutation prevents it. Inversion and duplication may provide an additional route to parity (8), and other biological explanations have been proposed (31); the simulation establishes the sufficiency and boundary condition of the mutation-based mechanism rather than excluding those contributions.

---

**Figure S1:**
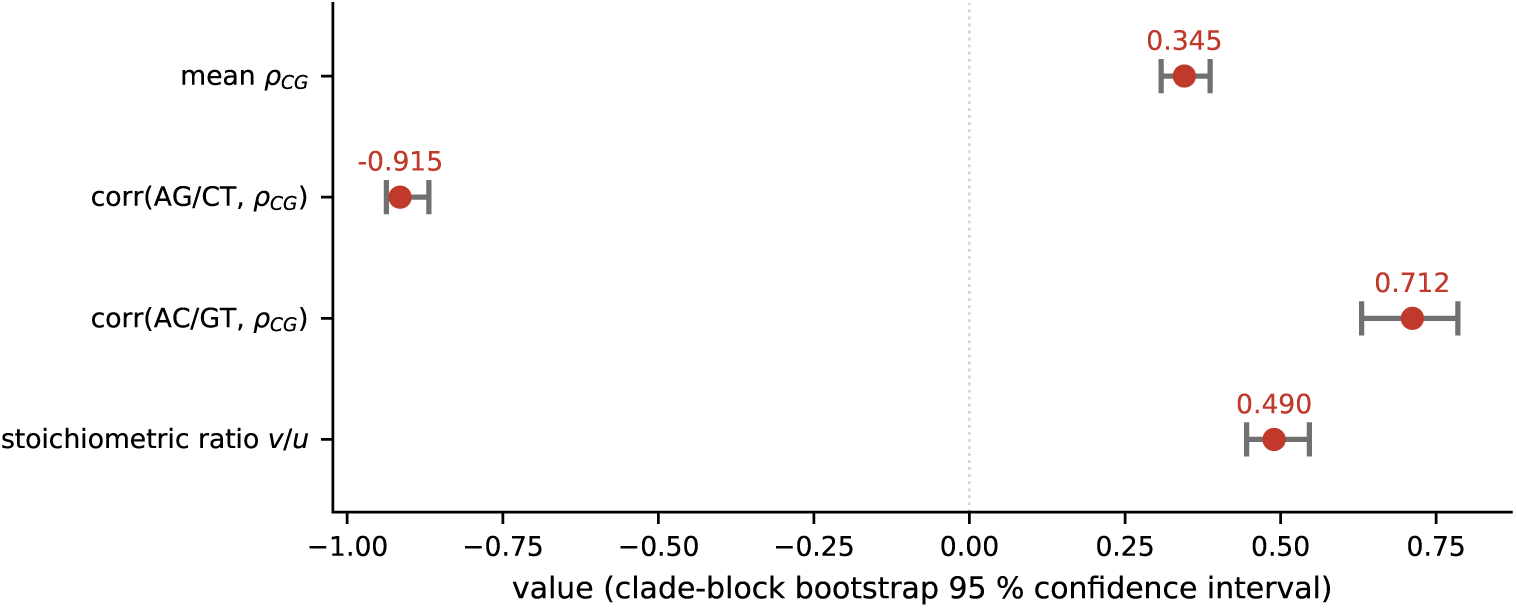
Principal statistics with clade-block bootstrap 95 % confidence intervals obtained by resampling taxonomic orders. Red points are point estimates and grey intervals are confidence intervals. Mean *ρ*_CG_ quantifies CpG depletion; the correlations describe the AG/CT and AC/GT associations with *ρ*_CG_; and *v/u* is the TpG/CpA-to-CpG mass-balance statistic, whose theoretical expectation is 0.5 (Supplementary Table S1).

**Figure S2:**
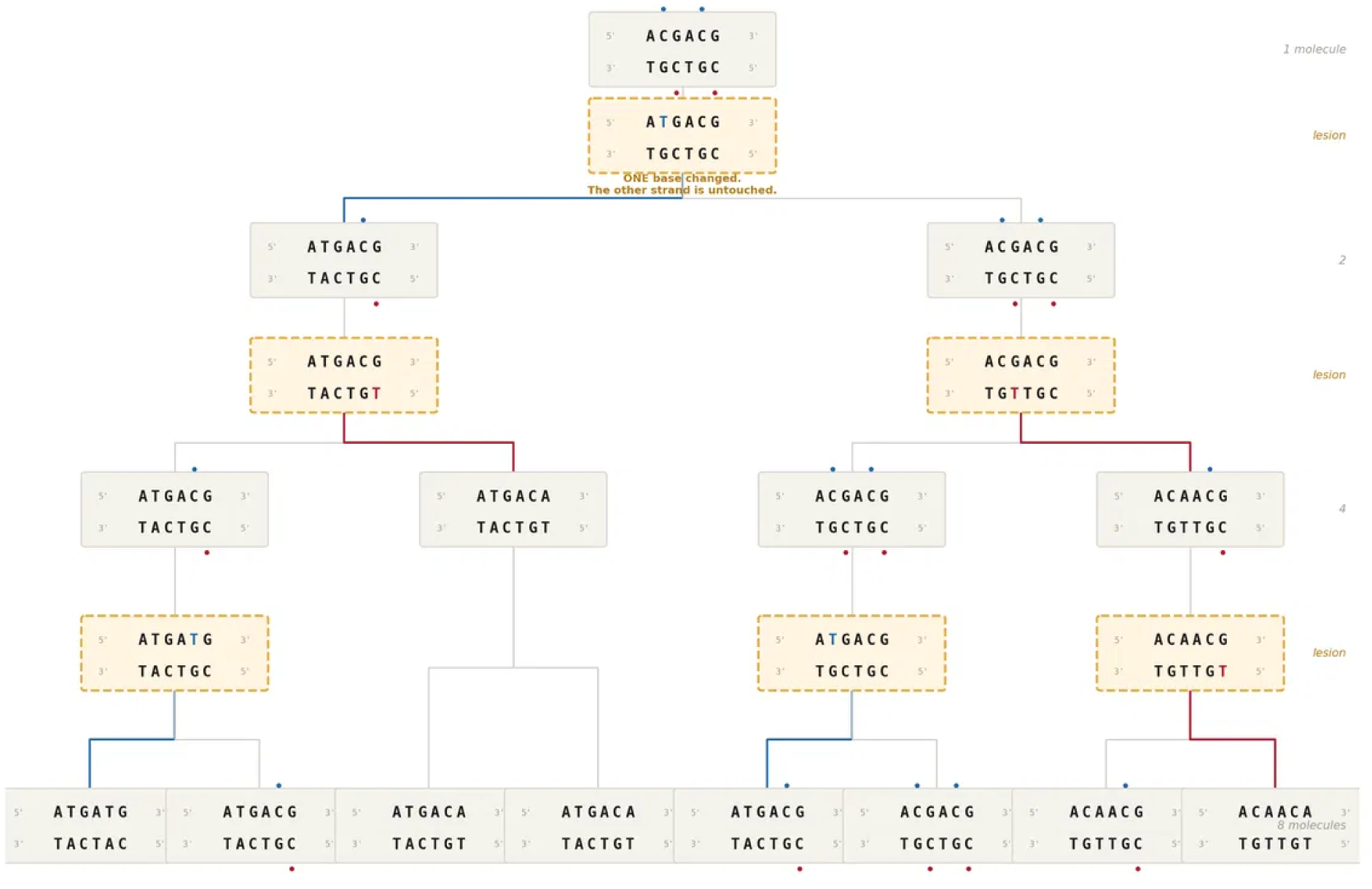
Fixation of methyl-CpG deamination through replication. A deamination event alters one cytosine on one strand and leaves the opposite strand unchanged, creating a mismatch. The daughter that inherits the altered parental strand fixes TpG and its reverse-complement representation CpA; the daughter inheriting the unaltered strand reconstructs CpG. Independent events in either orientation generate both products across descendant molecules. Blue and red marks identify the two parental-strand orientations.

**Figure S3:**
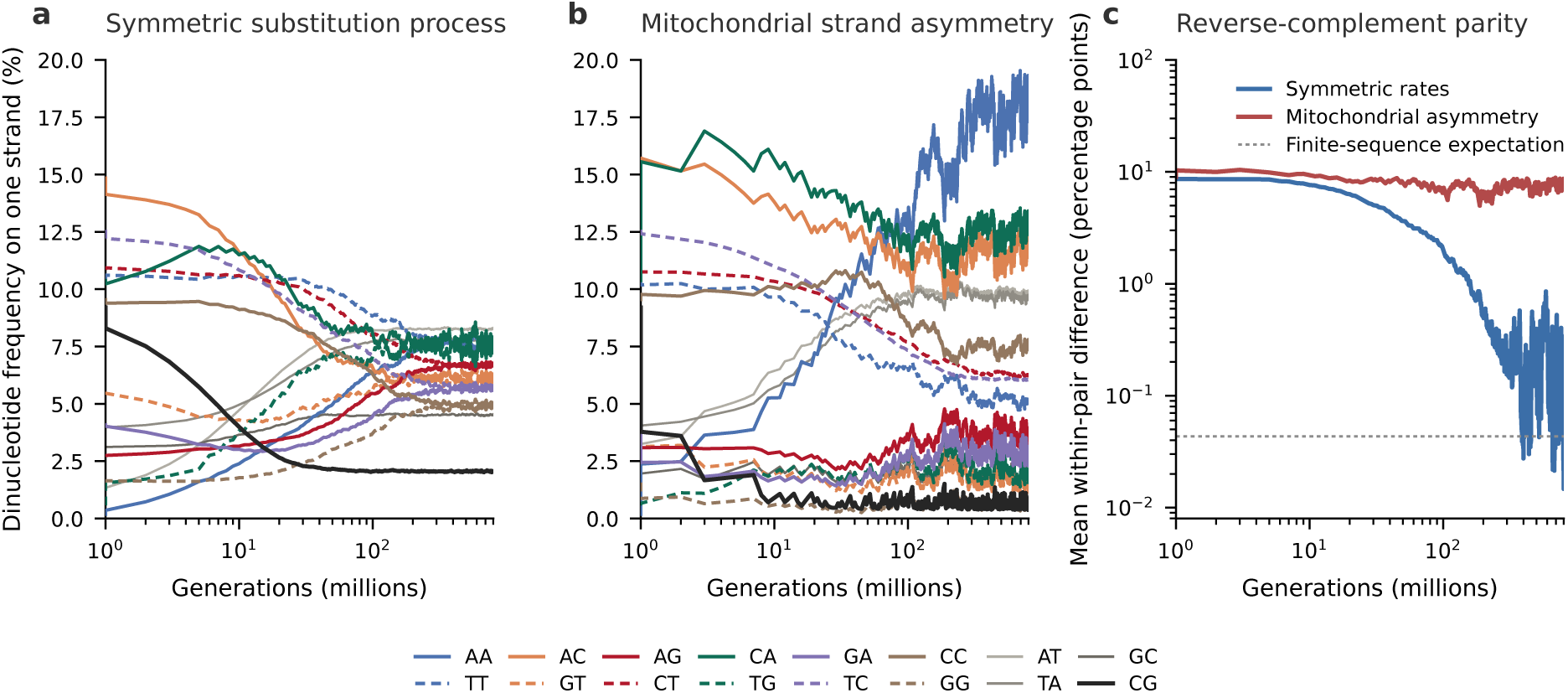
Reverse-complement parity under symmetric and strand-biased mutation. Both DNA strands are represented explicitly, and both runs start from the same sequence designed to violate parity in all six reverse-complement pairs. **(A)** Under the same context-dependent rates on both strands, the two members of every pair converge. Within each colour, solid and dashed lines represent the two members of one pair; AT, TA, CG and GC are self-reverse-complementary and are shown separately. **(B)** With the measured human mitochondrial strand biases—nine-fold for G-to-A and 1.8-fold for T-to-C on the light strand relative to the heavy strand (30)—the paired frequencies do not converge. **(C)** Mean absolute within-pair frequency difference on a logarithmic scale. The symmetric regime reaches the finite-sequence sampling floor, whereas the strand-biased regime remains above it.

**Figure S4:**
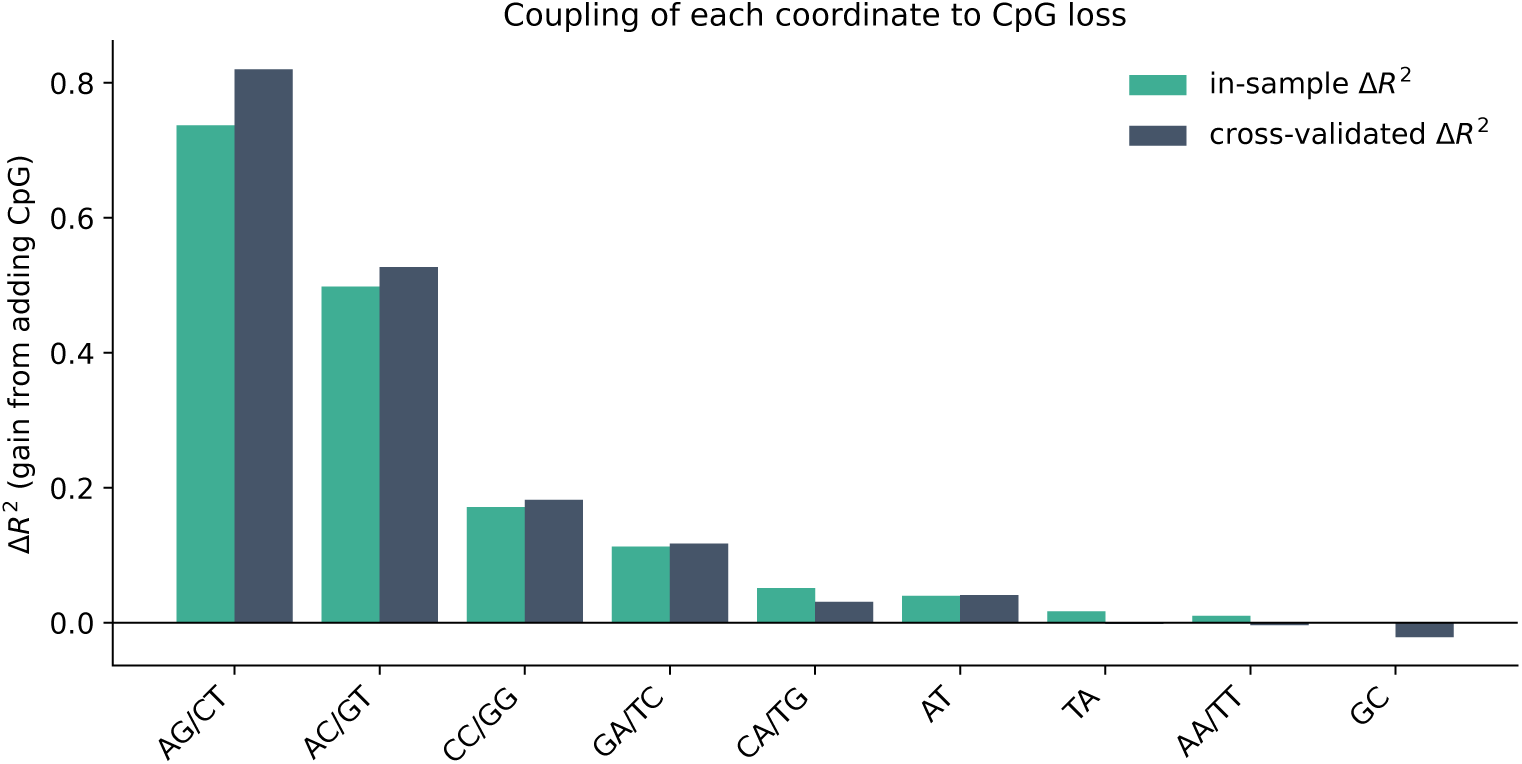
Gain in explained variance (Δ*R*^2^) from adding the CpG frequency to a base-composition model, for each strand-symmetric coordinate, in sample and under leave-one-order-out cross-validation. AG/CT has the largest gain both in and out of sample (Supplementary Table S2).

**Figure S5:**
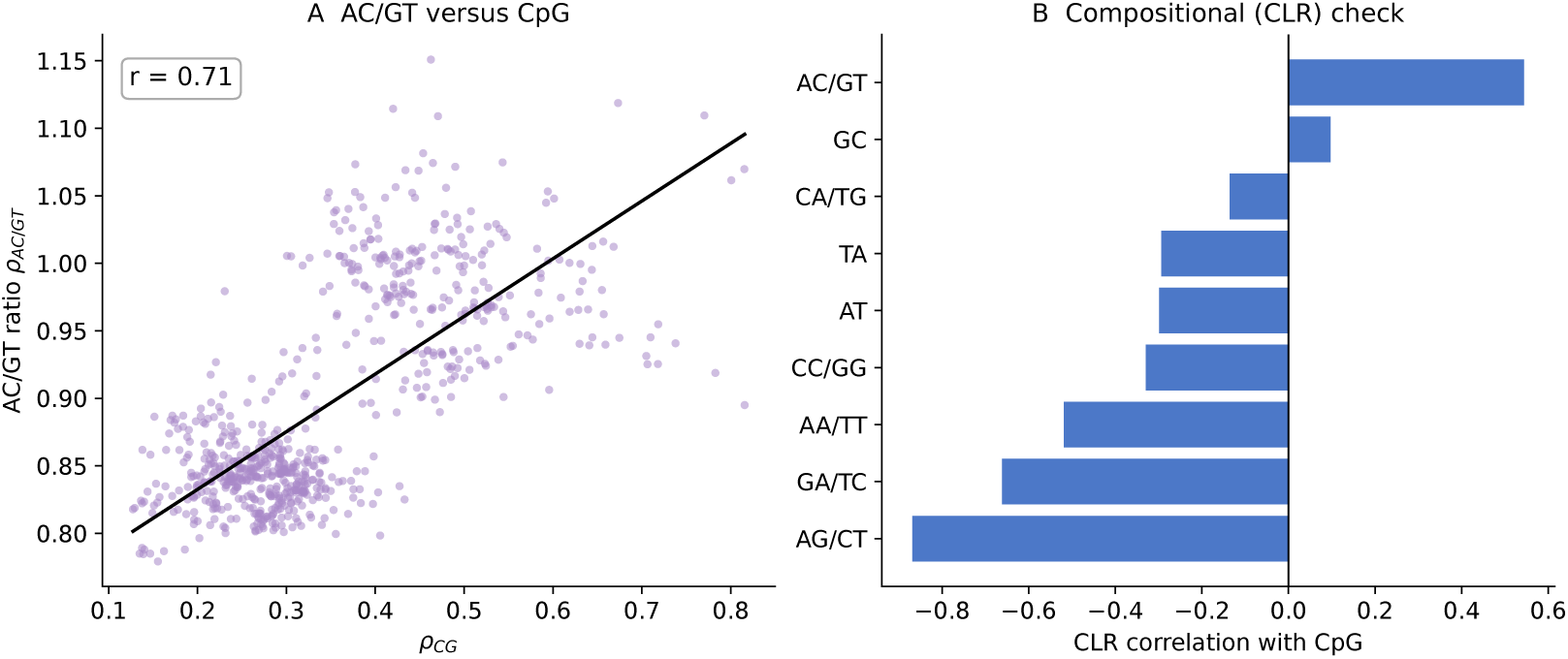
**(A)** AC/GT ratio against *ρ*_CG_ across the 753 vertebrate genomes. **(B)** Correlation of each coordinate with CpG after centred-log-ratio transformation of the complete sixteen-part dinucleotide composition. AG/CT remains the strongest association under this compositional treatment (Supplementary Table S3).

**Figure S6:**
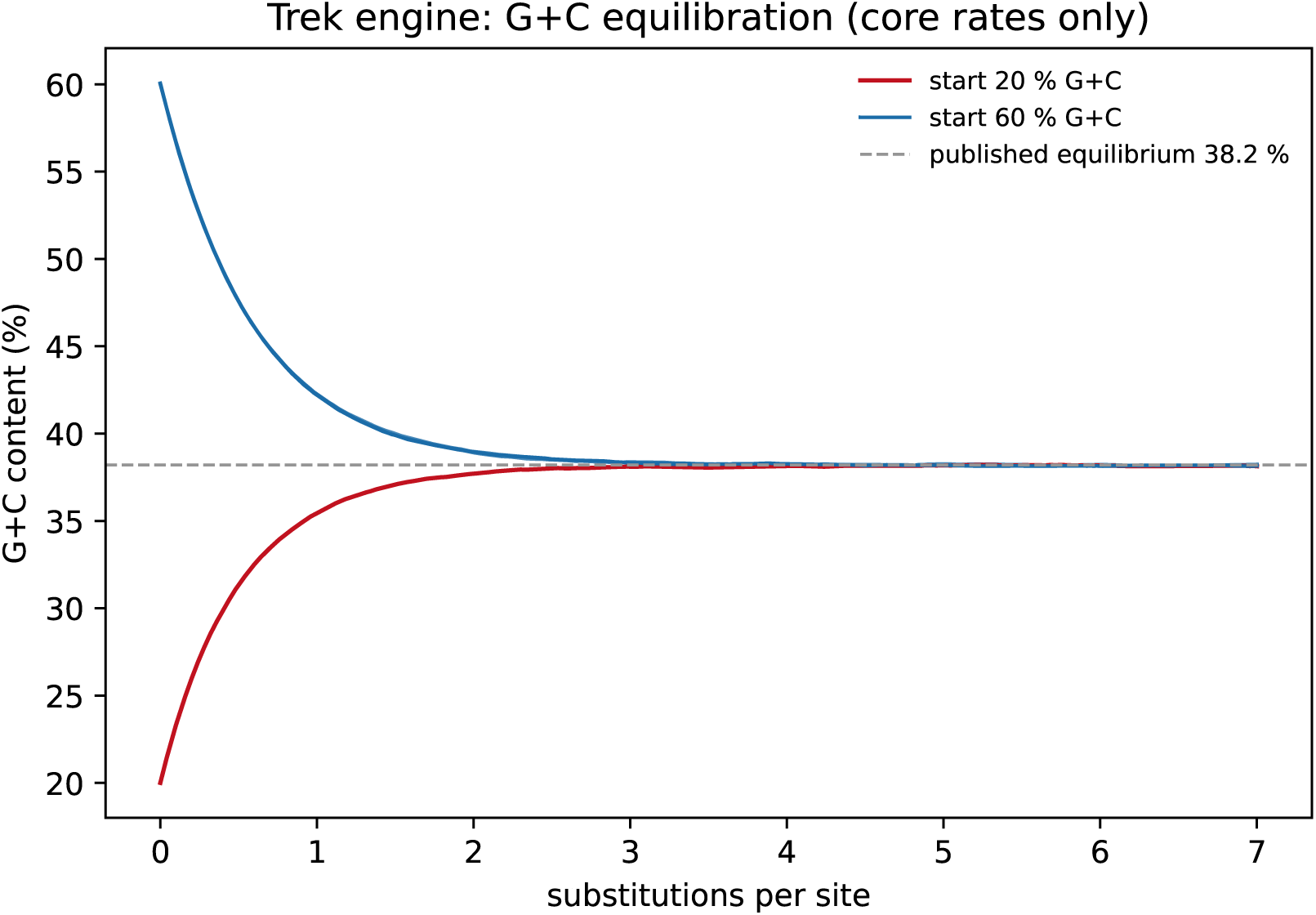
Validation of the forward-evolution engine. Under the Trek core seven-mer germline rates, sequences with high and low starting G+C converge to 38.2 % G+C, reproducing the published equilibrium A/T/G/C composition of 30.9*/*30.9*/*19.1*/*19.1 %. The core rates exclude the elevated mutability of methylated CpG.

**Figure S7:**
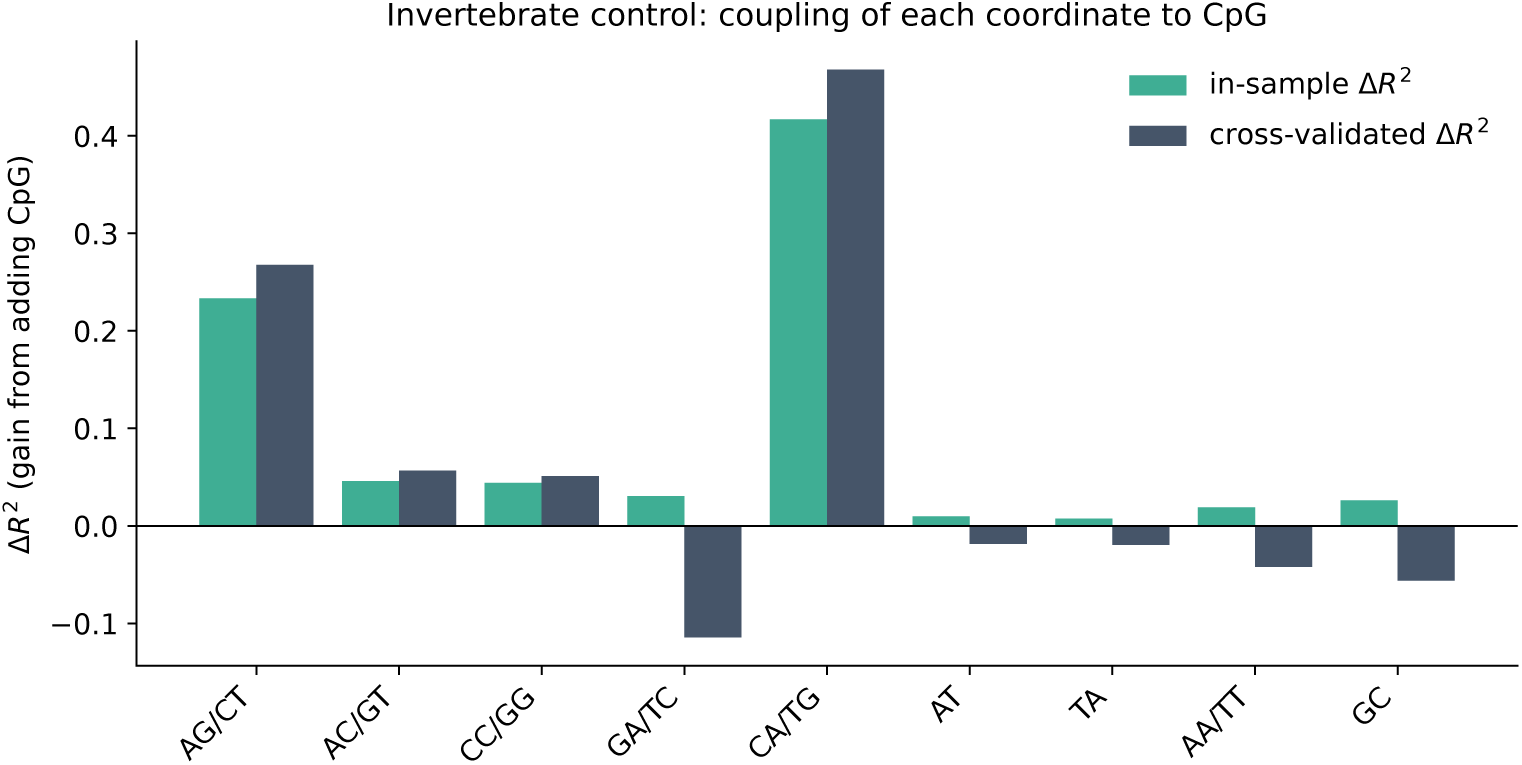
CpG-coupling analysis applied to the 481 invertebrate genomes. AG/CT coupling is substan- tially weaker than in vertebrates, and the largest invertebrate gain occurs for CA/TG rather than AG/CT (Supplementary Table S9).

**Figure S8:**
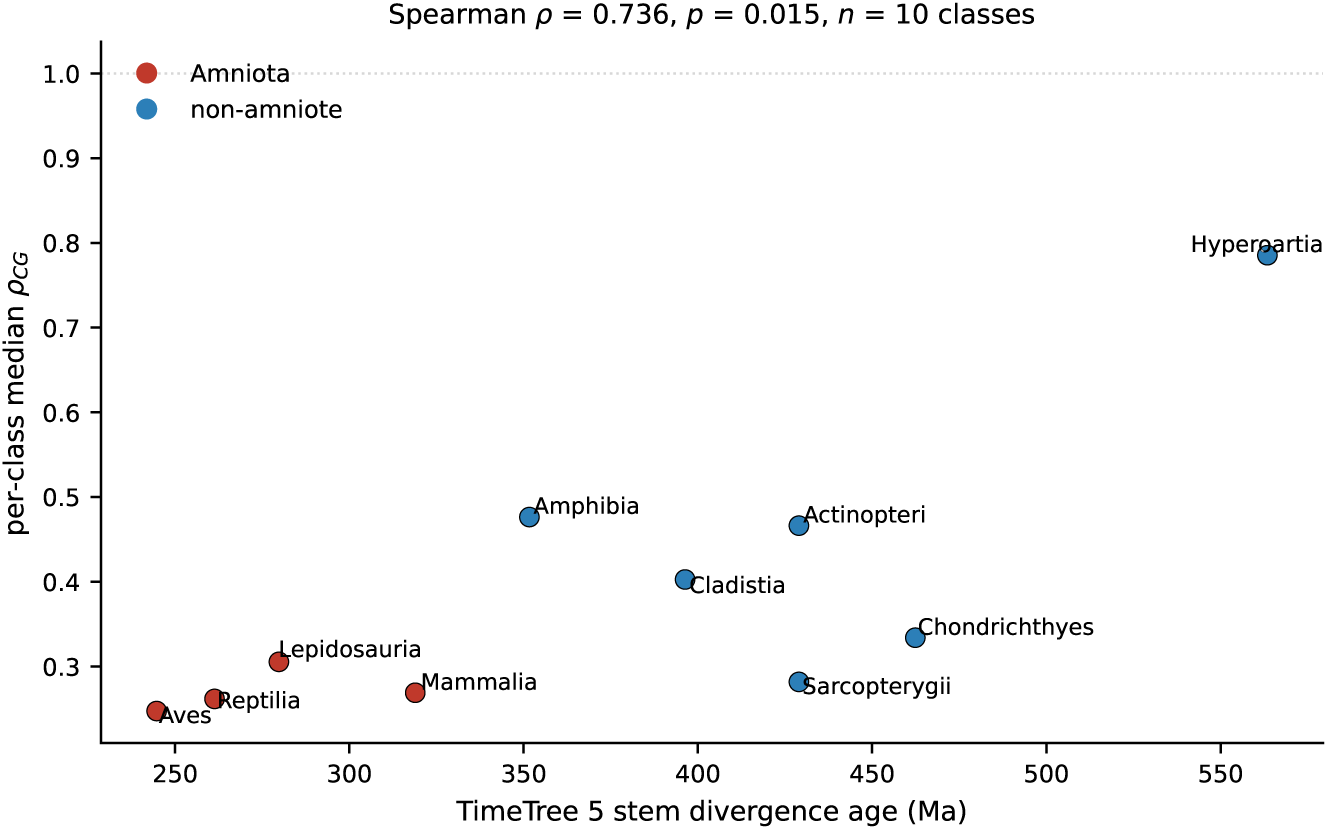
Class median *ρ*_CG_ against TimeTree 5 stem divergence age (24). The positive association denotes weaker CpG depletion at older stem ages. It is based on ten non-independent class values and is sensitive to Hyperoartia (Supplementary Table S10).

**Figure S9:**
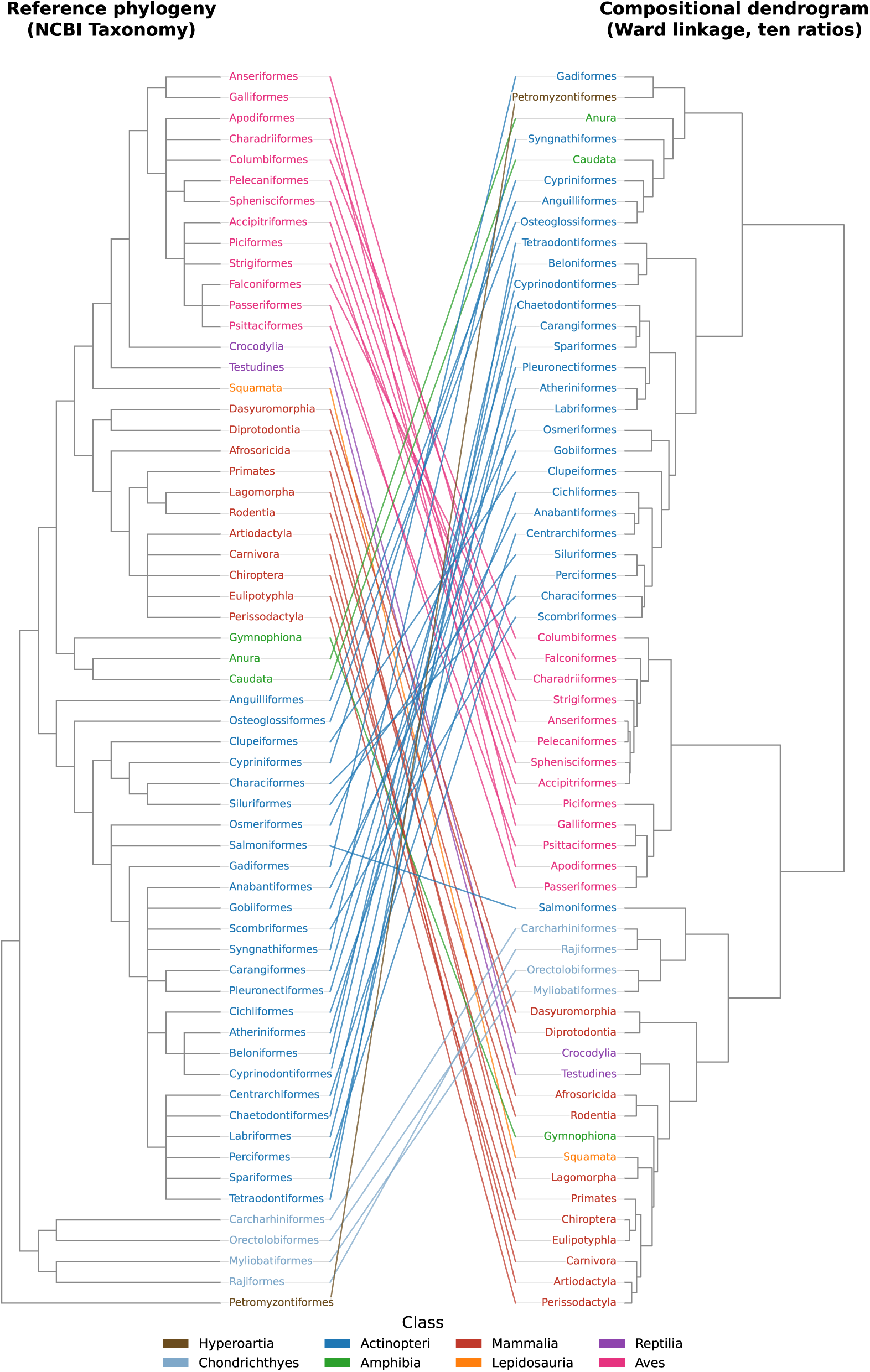
Tanglegram comparing the NCBI Taxonomy vertebrate topology (25) (left) with an un- supervised Ward dendrogram of the ten ratios (right) for the 60 orders represented by at least three genomes. Matching orders are connected by class-coloured lines. Related orders often remain close, whereas shared CpG depletion draws many amniote orders together in the compositional tree. Cophenetic-distance agreement is substantial but incomplete (Mantel *r* = 0.735, *p <* 0.001).

**Figure S10:**
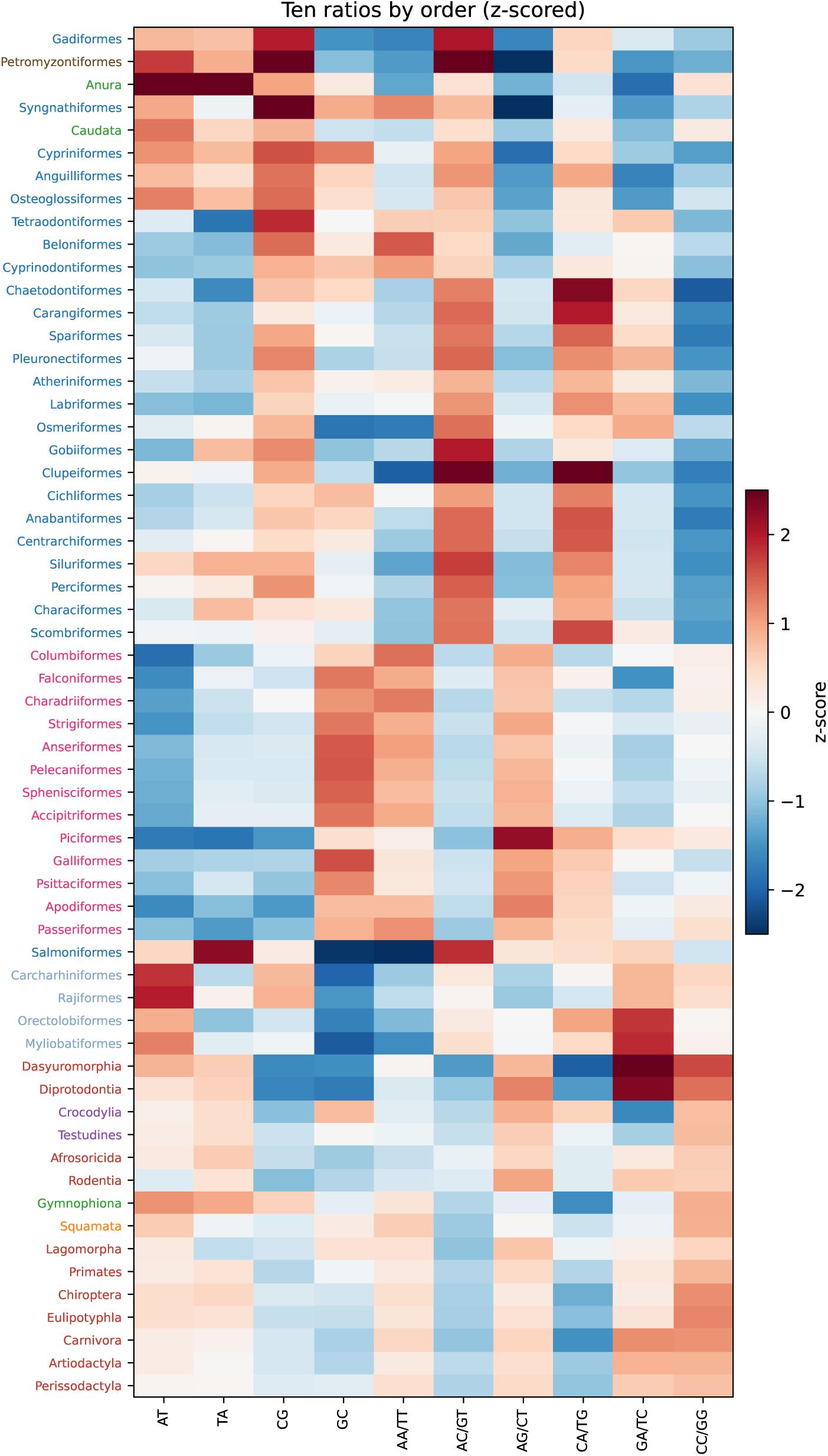
Heatmap of the ten z-scored strand-symmetric ratios by vertebrate order. Orders follow the Ward dendrogram in Supplementary Figure S9, and labels are coloured by class. Blocks of similar CpG depletion and associated coordinate values extend across several amniote orders.

**Figure S11:**
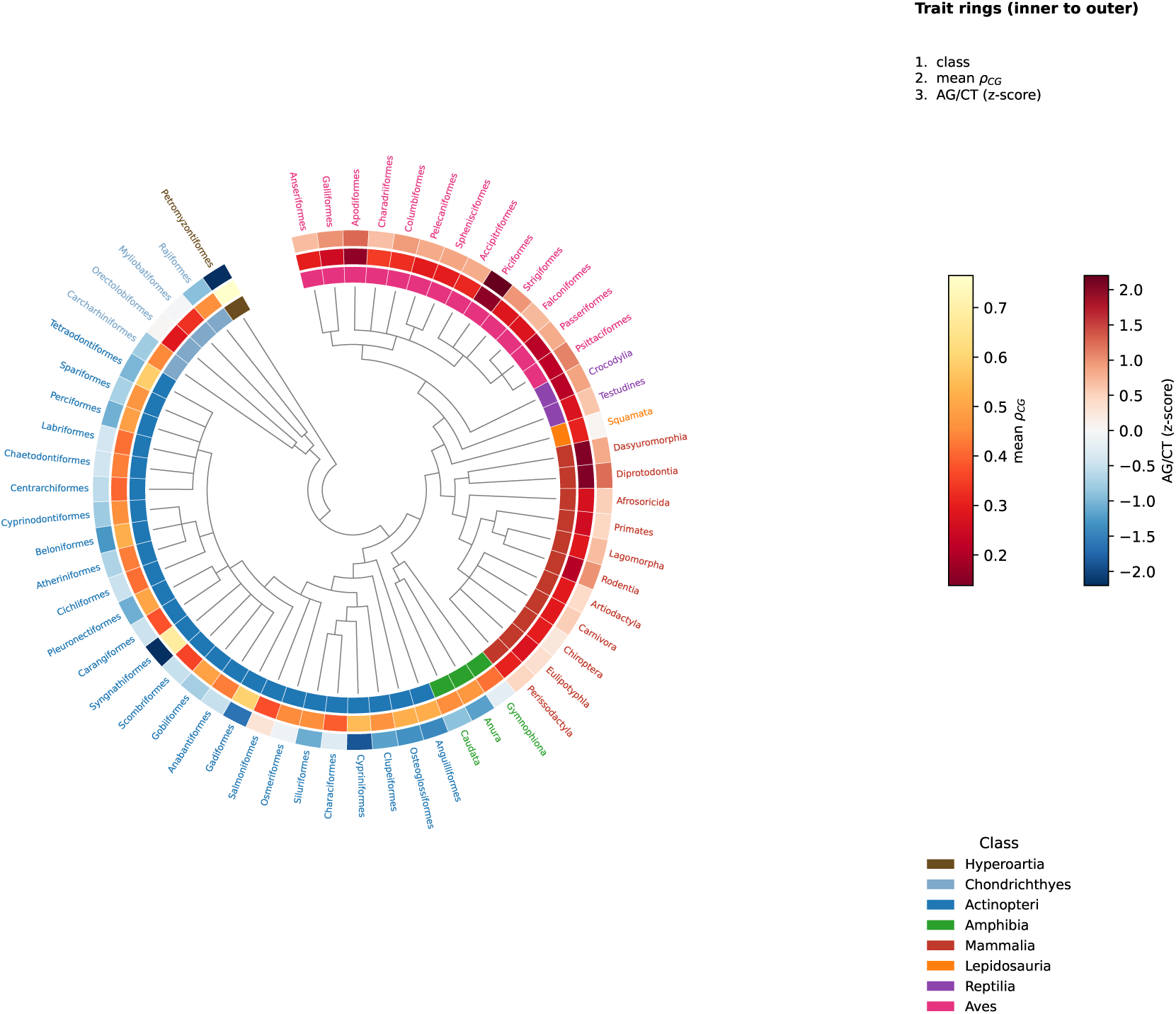
Radial rendering of 60 vertebrate orders. The inner circle shows the NCBI phylogenetic tree (25), the middle circle shows the mean CpG ratio *ρ*_CG_, and the outer circle shows the z-scored AG/CT ratio. The CpG-depletion gradient and associated AG/CT increase are concentrated in amniote lineages but also vary across the broader tree.

**Figure S12:**
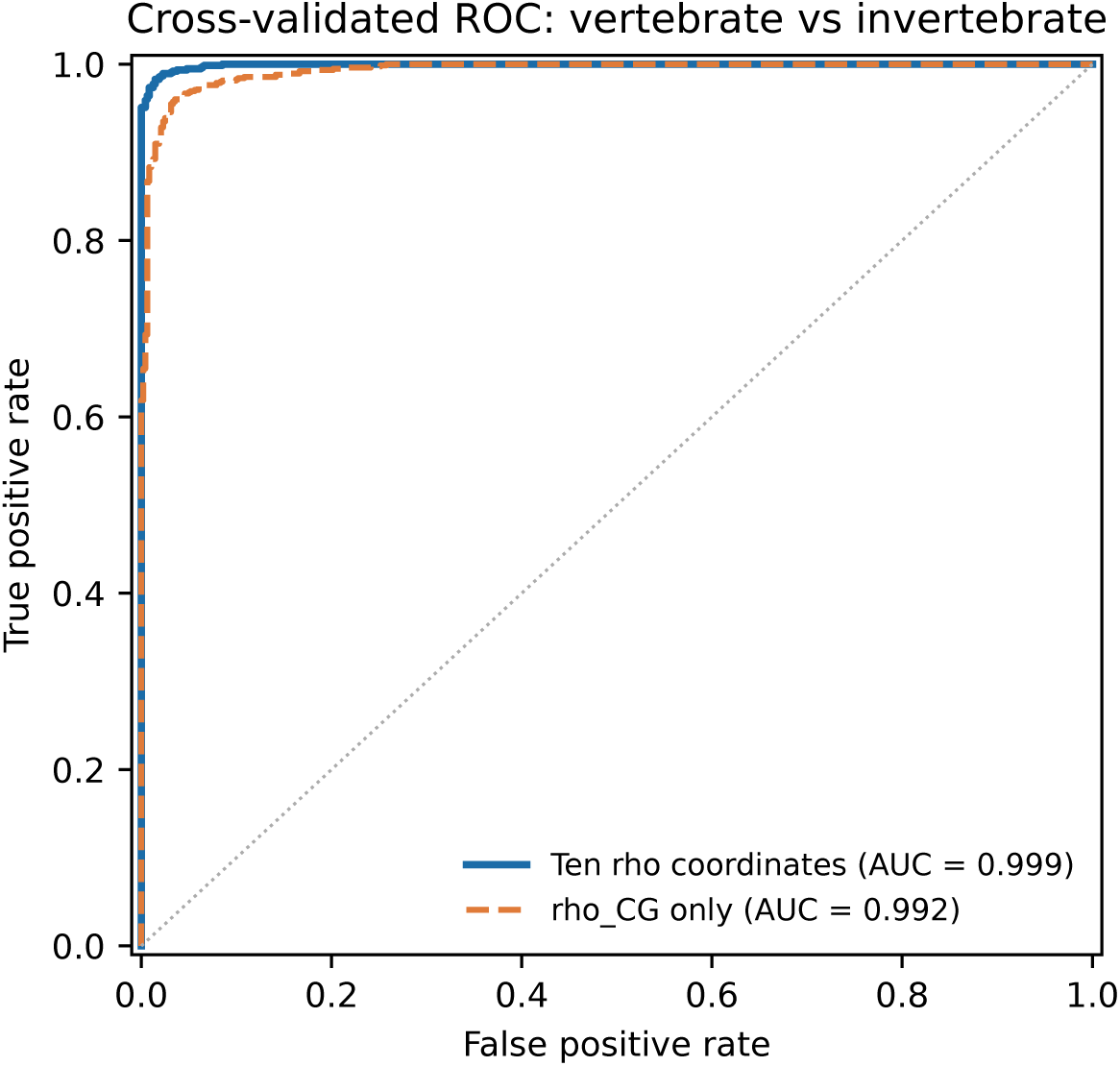
Pooled out-of-fold receiver operating characteristic curve for the class-balanced random forest separating vertebrates from invertebrates using the ten ratios, under stratified ten-fold cross-validation (AUC = 0.999).

**Figure S13:**
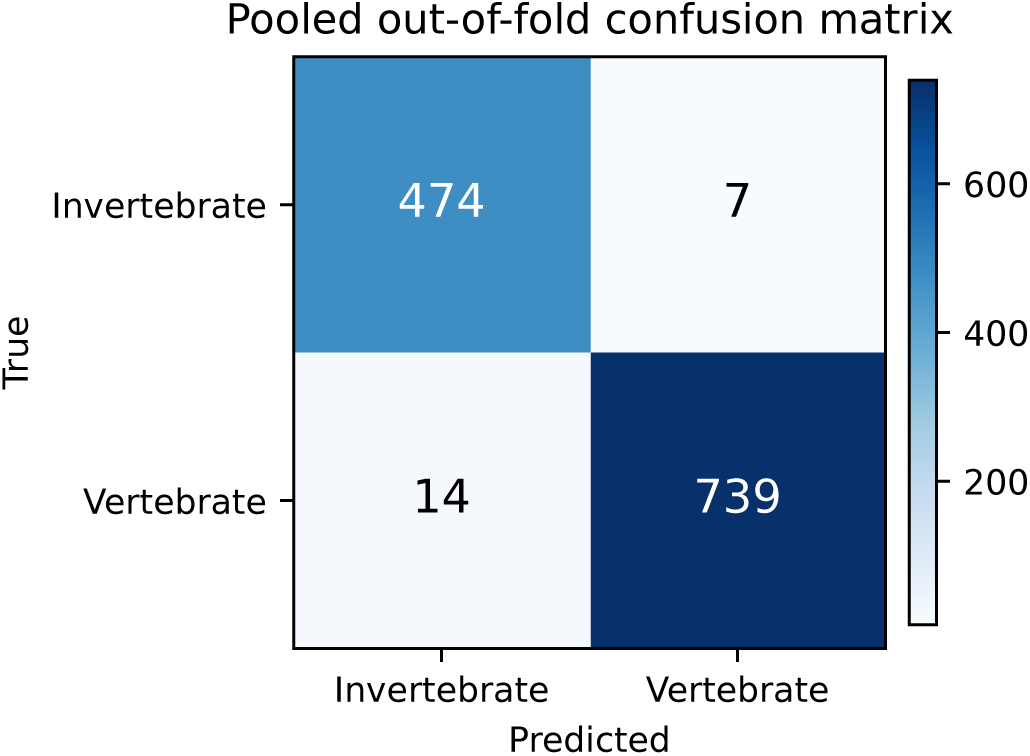
Confusion matrix from pooled out-of-fold predictions. The 14 false-negative vertebrates include comparatively weakly depleted pipefishes, seahorses and lampreys; the 7 false-positive invertebrates are unusually CpG-poor.

**Figure S14:**
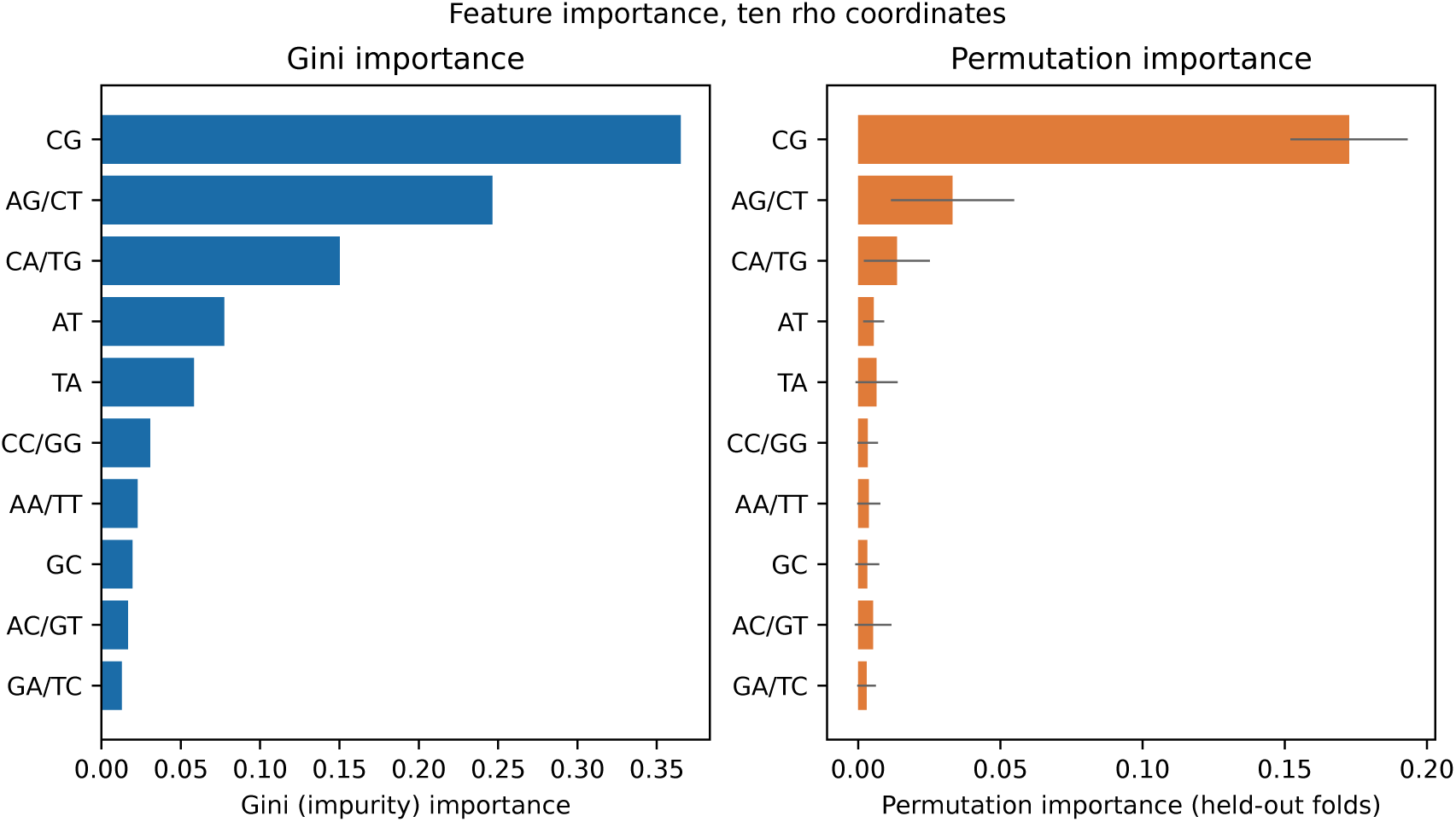
Random-forest feature importance estimated by impurity reduction and held-out permutation. Both methods rank *ρ*_CG_ first and AG/CT second; permutation importance is evaluated on held-out rather than training observations.

**Figure S15:**
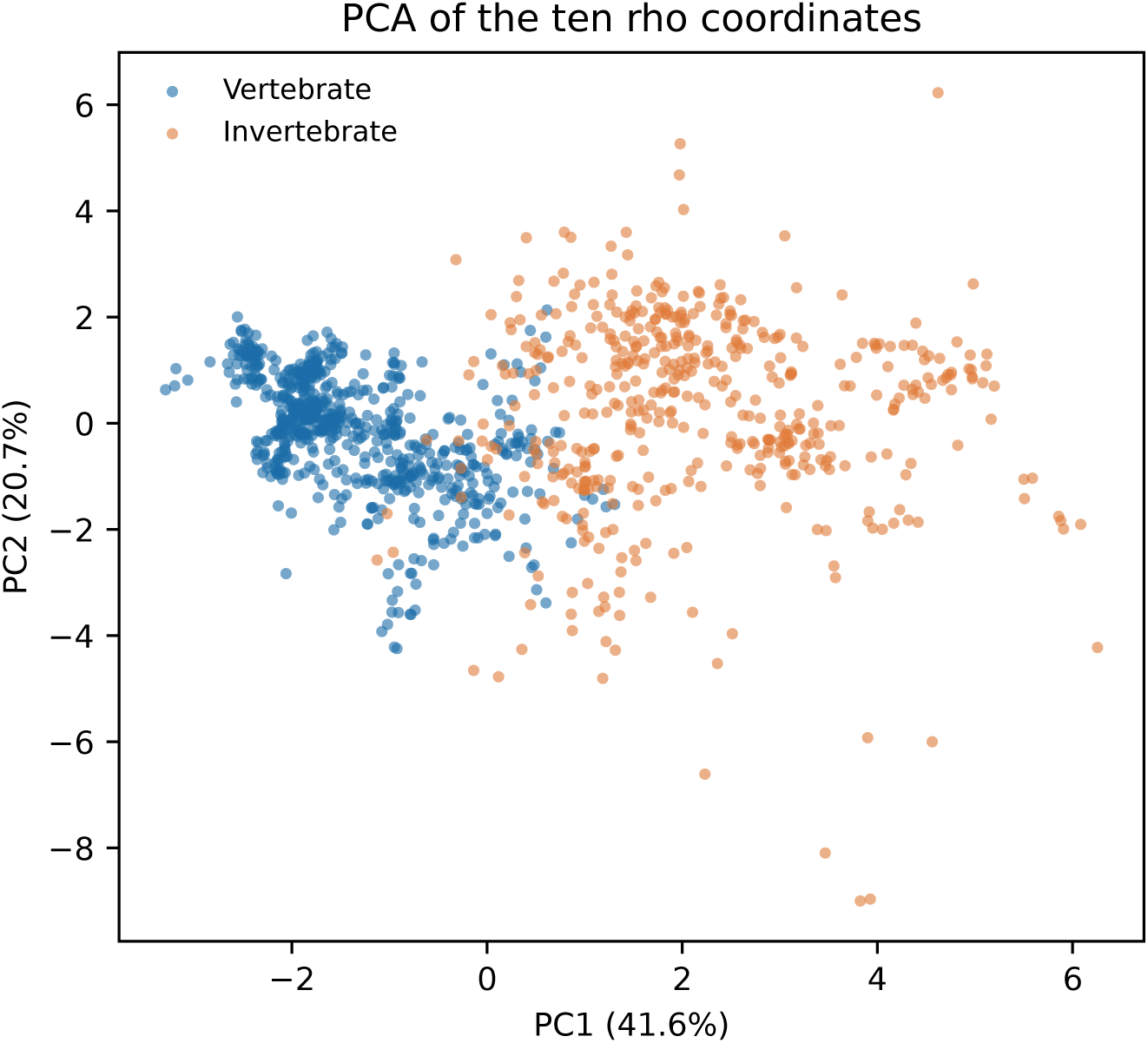
Principal component analysis of the ten standardised ratios, coloured by animal group. PC1 and PC2 explain 41.6 % and 20.7 % of the variance, respectively. CpG and AG/CT have opposite PC1 loadings along the principal direction separating the groups.

**Table S1:** Principal genome-level statistics with clade-block bootstrap 95 % confidence intervals from 2000 resamples of taxonomic orders. Mean *ρ*_CG_ quantifies CpG depletion; the two correlations measure the direction and strength of the AG/CT and AC/GT associations with *ρ*_CG_; and *v/u* is the ratio of mean TpG/CpA excess to mean CpG deficit, with theoretical expectation 0.5. The statistics are visualised in Supplementary Figure S1.

| Statistic | Point estimate | 95 % CI |
| --- | --- | --- |
| mean $\rho_{CG}$ | 0.345 | [0.308, 0.387] |
| corr(AG/CT, $\rho_{CG}$) | -0.915 | [-0.937, -0.869] |
| corr(AC/GT, $\rho_{CG}$) | 0.712 | [0.630, 0.785] |
| stoichiometric ratio $v/u$ | 0.490 | [0.445, 0.546] |

**Table S2:** Coupling of each non-CG strand-symmetric coordinate to CpG across the 753 vertebrate genomes. “Base only” and “Base + CpG” give the in-sample *R*^2^ of the nested models; Δ*R*^2^ is the leave-one-order-out cross-validated gain. ΔAIC and ΔBIC compare the augmented model with the baseline, and *p_adj_* is the Benjamini–Hochberg-adjusted significance of the CpG term. See Supplementary Figure S4.

| Coord. | Base only | Base+CpG | $\Delta R^2$ | $\Delta R^2_{cv}$ | $\Delta AIC$ / $\Delta BIC$ | $p_{adj}$ |
| --- | --- | --- | --- | --- | --- | --- |
| AG/CT | 0.152 | 0.889 | 0.737 | 0.820 | -1529 / -1524 | $< 10^{-300}$ |
| AC/GT | 0.003 | 0.501 | 0.498 | 0.527 | -520 / -515 | $3.0 \times 10^{-114}$ |
| CC/GG | 0.606 | 0.778 | 0.172 | 0.182 | -429 / -424 | $8.4 \times 10^{-95}$ |
| GA/TC | 0.073 | 0.186 | 0.113 | 0.117 | -96 / -91 | $1.1 \times 10^{-22}$ |
| CA/TG | 0.091 | 0.142 | 0.051 | 0.031 | -42 / -37 | $6.3 \times 10^{-11}$ |
| AT | 0.727 | 0.767 | 0.040 | 0.041 | -117 / -113 | $2.9 \times 10^{-27}$ |
| TA | 0.605 | 0.622 | 0.017 | -0.002 | -31 / -26 | $1.2 \times 10^{-8}$ |
| AA/TT | 0.777 | 0.787 | 0.010 | -0.003 | -33 / -28 | $4.9 \times 10^{-9}$ |
| GC | 0.550 | 0.550 | 0.000 | -0.021 | 1 / 6 | 0.437 |

**Table S3:** Robustness of selected associations with *ρ*_CG_ to taxonomic non-independence. Pearson correlations were calculated across genomes and after aggregation by order and class. A taxonomy- structured generalised least-squares (GLS) model then estimated the slope significance and adjusted correlation. AG/CT remains strongly associated at every level, whereas the AC/GT association disappears after GLS adjustment. See Supplementary Figure S5.

| Coord. | genome $r$ | order-mean $r$ | class-mean $r$ | GLS $p$ | GLS $r_{phylo}$ |
| --- | --- | --- | --- | --- | --- |
| AG/CT | -0.915 | -0.863 | -0.953 | $5.9 \times 10^{-64}$ | -0.562 |
| AC/GT | 0.712 | 0.731 | 0.908 | 0.878 | 0.006 |
| CA/TG | 0.266 | 0.404 | 0.390 | $1.9 \times 10^{-47}$ | -0.493 |

**Table S4:**
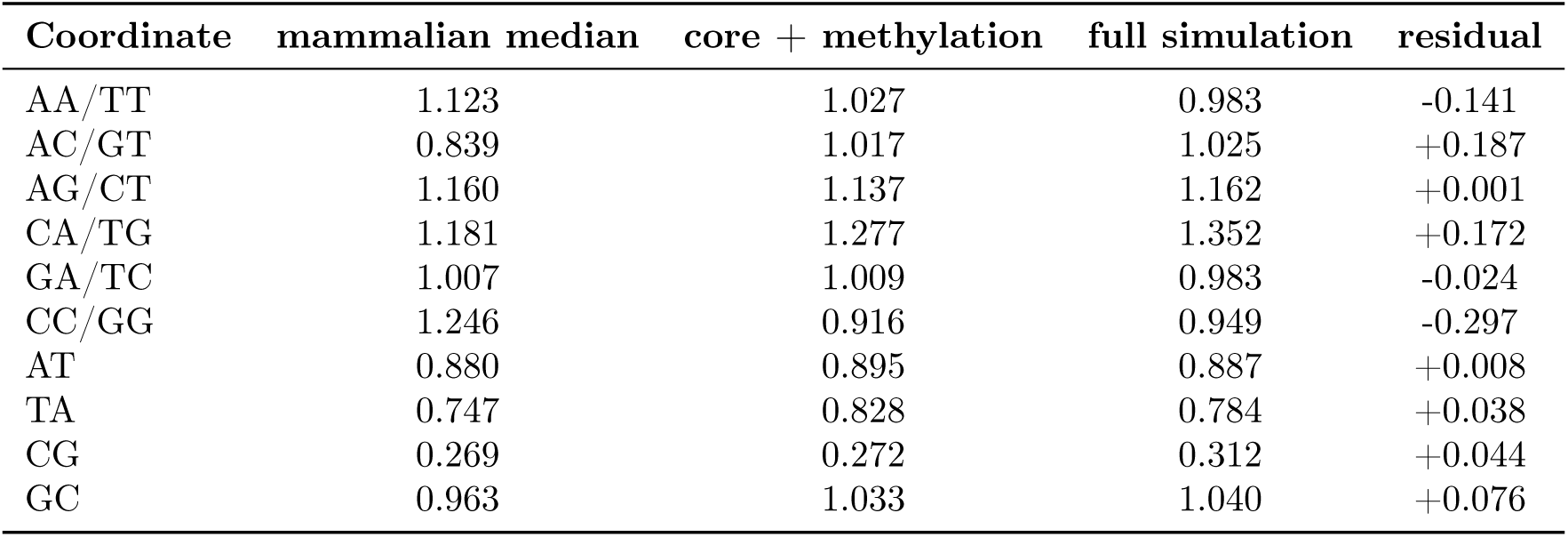
Observed mammalian median and simulated *ρ* values for the ten strand-symmetric coordinates. “Core + methylation” includes Trek substitution rates and the calibrated CpG term; the full simulation additionally includes the gBGC approximation. Residuals are full-simulation values minus mammalian medians. No replication-slippage parameter was fitted.

| Coordinate | mammalian median | core + methylation | full simulation | residual |
| --- | --- | --- | --- | --- |
| AA/TT | 1.123 | 1.027 | 0.983 | -0.141 |
| AC/GT | 0.839 | 1.017 | 1.025 | +0.187 |
| AG/CT | 1.160 | 1.137 | 1.162 | +0.001 |
| CA/TG | 1.181 | 1.277 | 1.352 | +0.172 |
| GA/TC | 1.007 | 1.009 | 0.983 | -0.024 |
| CC/GG | 1.246 | 0.916 | 0.949 | -0.297 |
| AT | 0.880 | 0.895 | 0.887 | +0.008 |
| TA | 0.747 | 0.828 | 0.784 | +0.038 |
| CG | 0.269 | 0.272 | 0.312 | +0.044 |
| GC | 0.963 | 1.033 | 1.040 | +0.076 |

**Table S5:** Per-genome transposable-element analysis from archived RepeatMasker annotations. TE fraction is the proportion assigned to the included element classes. Genome-wide, within-element, non-element and *>* 5 kb *ρ*_CG_ values are calculated with the corresponding region’s base composition. Flanking bins are defined by base-wise distance from the nearest element: 0–100, 100–500, 500–1000, 1000–2000, 2000–5000 and *>* 5000 bp; non-element repeat classes remain in the flanks. Genomes with TE fraction below 2 % were excluded from pooled tests. Across the 11 retained genomes, inside-element and *>* 5 kb values differ by a two-sided exact Wilcoxon signed-rank test (*p* = 0.0098).

| Organism | Class | TE frac. | $\rho_{CG}$ | in TE | non-TE | $> 5$ kb | used |
| --- | --- | --- | --- | --- | --- | --- | --- |
| *Crocodylus porosus* | Reptilia | 0.361 | 0.199 | 0.184 | 0.209 | 0.465 | yes |
| *Gavialis gangeticus* | Reptilia | 0.360 | 0.195 | 0.186 | 0.201 | 0.427 | yes |
| *Rousettus aegyptiacus* | Mammalia | 0.313 | 0.344 | 0.248 | 0.380 | 0.623 | yes |
| *Xenopus tropicalis* | Amphibia | 0.312 | 0.334 | 0.462 | 0.268 | 0.281 | yes |
| *Miniopterus natalensis* | Mammalia | 0.271 | 0.287 | 0.202 | 0.313 | 0.459 | yes |
| *Condylura cristata* | Mammalia | 0.174 | 0.266 | 0.161 | 0.282 | 0.557 | yes |
| *Trachemys scripta* | Reptilia | 0.140 | 0.276 | 0.242 | 0.282 | 0.370 | yes |
| *Coturnix japonica* | Aves | 0.060 | 0.230 | 0.122 | 0.238 | 0.288 | yes |
| *Parus major* | Aves | 0.045 | 0.179 | 0.085 | 0.185 | 0.214 | yes |
| *Nothoprocta perdicaria* | Aves | 0.037 | 0.328 | 0.179 | 0.335 | 0.387 | yes |
| *Lacerta agilis* | Lepidosauria | 0.034 | 0.322 | 0.200 | 0.327 | 0.330 | yes |
| *Python bivittatus* | Lepidosauria | 0.012 | 0.217 | 0.120 | 0.219 | 0.225 | no |
| *Thamnophis sirtalis* | Lepidosauria | 0.002 | 0.303 | 0.307 | 0.303 | 0.303 | no |
| *Cyclopterus lumpus* | Actinopteri | 0.001 | 0.545 | 0.570 | 0.545 | 0.546 | no |
| *Anabas testudineus* | Actinopteri | 0.001 | 0.360 | 0.270 | 0.360 | 0.361 | no |
| *Electrophorus electricus* | Actinopteri | 0.000 | 0.411 | 0.155 | 0.411 | 0.412 | no |
| *Nanorana parkeri* | Amphibia | 0.000 | 0.434 | 0.412 | 0.434 | 0.434 | no |

**Table S6:**
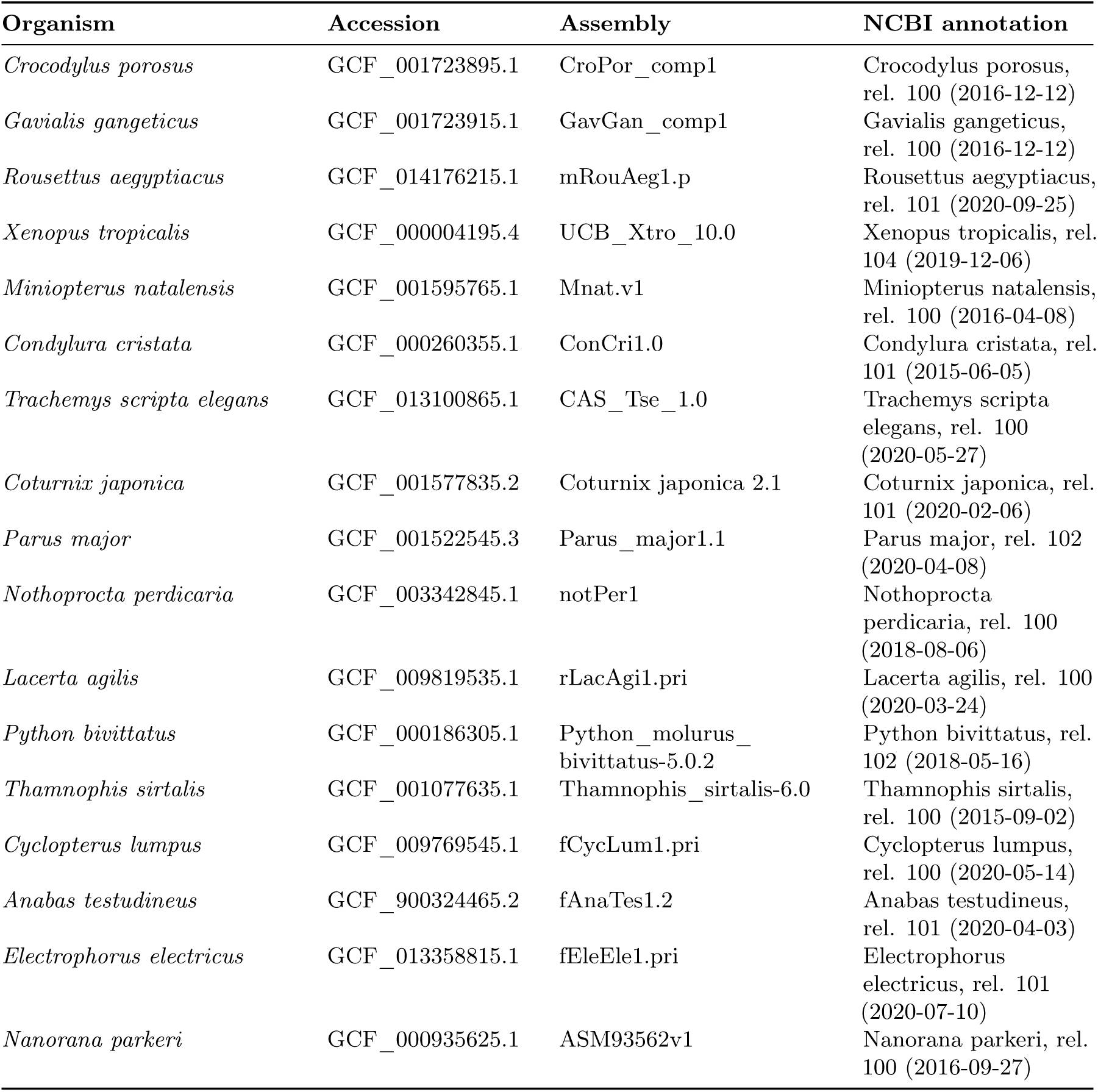
Assembly and annotation provenance for the seventeen genomes in Supplementary Table S5, shown in the same order. The versioned accession and NCBI annotation release identify the genome sequence and RepeatMasker output. FASTA (<accession>_genomic.fna.gz) and annotation (<accession>_rm.out.gz) files were obtained from the NCBI RefSeq FTP tree. Provenance for the sixteen isochore genomes is recorded in results/00_data_prep/genome_provenance.csv.

**Table S7:** Separation of two candidate vertebrate groupings. Silhouette coefficient and cross-validated logistic-regression accuracy were calculated from the ten standardised ratios; PERMANOVA used Euclidean distances in MFA space and 4999 permutations. Each measure favours the amniote–non-amniote partition. Median *ρ*_CG_ is 0.291 in ectothermic amniotes, 0.264 in endothermic amniotes and approximately 0.44 in ectothermic non-amniotes.

| Grouping | silhouette | CV accuracy | PERMANOVA $R^2$ | pseudo-$F$ |
| --- | --- | --- | --- | --- |
| amniote / non-amniote | 0.371 | 0.993 | 0.260 | 263.3 |
| endothermic / ectothermic | 0.284 | 0.954 | 0.211 | 200.7 |

**Table S8:** Isochore coverage and median *ρ*_CG_ by Bernardi family in ten amniote and six non-amniote genomes. Coverage is the percentage of genome sequence assigned to each family. H3 coverage is 3.0-fold greater in amniotes (Mann–Whitney *p* = 0.022). Median *ρ*_CG_ increases monotonically across families in all ten amniotes and in four of six non-amniotes; the gradient is therefore not exclusive to Amniota.

| Family | cover. amniote (%) | cover. non-amn. (%) | $\rho_{CG}$ amniote | $\rho_{CG}$ non-amn. |
| --- | --- | --- | --- | --- |
| L1 | 13.63 | 13.11 | 0.171 | 0.497 |
| L2 | 24.93 | 30.44 | 0.200 | 0.385 |
| H1 | 41.80 | 38.76 | 0.219 | 0.476 |
| H2 | 13.50 | 14.68 | 0.300 | 0.582 |
| H3 | 3.89 | 1.26 | 0.439 | 0.228 |

**Table S9:**
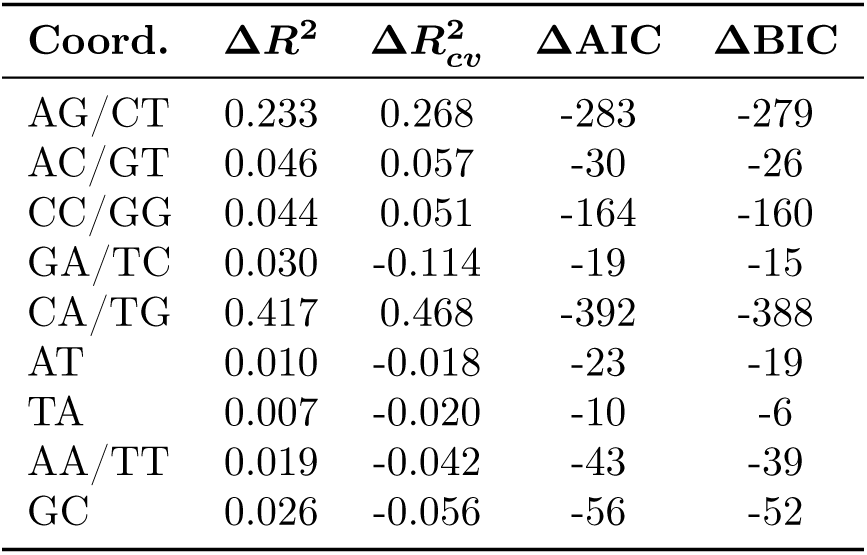
CpG-coupling analysis applied without modification to the 481 invertebrate genomes. AG/CT coupling is weaker than in vertebrates, and the largest invertebrate Δ*R*^2^ occurs for CA/TG. See Supplementary Figure S7.

| Coord. | $\Delta R^2$ | $\Delta R^2_{cv}$ | $\Delta AIC$ | $\Delta BIC$ |
| --- | --- | --- | --- | --- |
| AG/CT | 0.233 | 0.268 | -283 | -279 |
| AC/GT | 0.046 | 0.057 | -30 | -26 |
| CC/GG | 0.044 | 0.051 | -164 | -160 |
| GA/TC | 0.030 | -0.114 | -19 | -15 |
| CA/TG | 0.417 | 0.468 | -392 | -388 |
| AT | 0.010 | -0.018 | -23 | -19 |
| TA | 0.007 | -0.020 | -10 | -6 |
| AA/TT | 0.019 | -0.042 | -43 | -39 |
| GC | 0.026 | -0.056 | -56 | -52 |

**Table S10:** Class median *ρ*_CG_, sample size and TimeTree 5 stem divergence age. *ρ*_CG_ is positively associated with stem age (Spearman *ρ* = 0.736, *p* = 0.015), corresponding to weaker CpG depletion at older stem ages. The exploratory comparison contains ten non-independent class values and is sensitive to Hyperoartia. See Supplementary Figure S8.

| Class | $n$ | median $\rho_{CG}$ | TimeTree stem age (Ma) |
| --- | --- | --- | --- |
| Aves | 156 | 0.247 | 244.76 |
| Reptilia | 23 | 0.262 | 261.37 |
| Mammalia | 244 | 0.269 | 318.95 |
| Sarcopterygii | 2 | 0.282 | 429.00 |
| Lepidosauria | 31 | 0.305 | 279.83 |
| Chondrichthyes | 23 | 0.334 | 462.40 |
| Cladistia | 2 | 0.403 | 396.35 |
| Actinopteri | 235 | 0.466 | 429.00 |
| Amphibia | 33 | 0.476 | 351.69 |
| Hyperoartia | 4 | 0.785 | 563.39 |

**Table S11:**
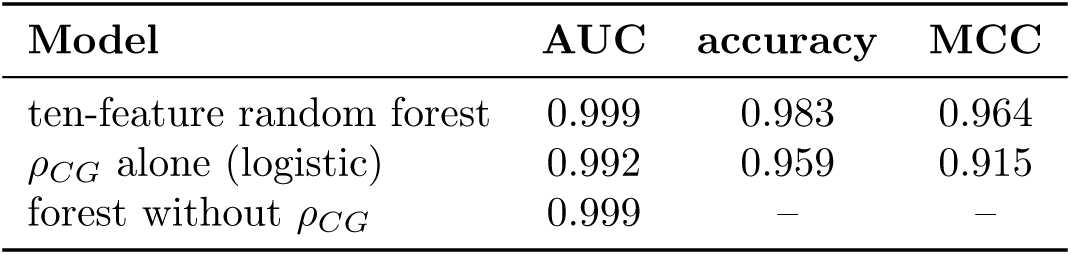
Vertebrate–invertebrate classification under stratified ten-fold cross-validation. The ten-feature random-forest confusion matrix is TN = 474, FP = 7, FN = 14 and TP = 739. *ρ*_CG_ alone approaches the full-model AUC, while the forest without *ρ*_CG_ retains the same reported AUC, demonstrating redundancy across the coordinated profile. See Supplementary Figures S12–S15.

